# Regulation of the MLH1-MLH3 endonuclease in meiosis

**DOI:** 10.1101/2020.02.12.946293

**Authors:** Elda Cannavo, Aurore Sanchez, Roopesh Anand, Lepakshi Ranjha, Jannik Hugener, Céline Adam, Ananya Acharya, Nicolas Weyland, Xavier Aran-Guiu, Jean-Baptiste Charbonnier, Eva R. Hoffmann, Valérie Borde, Joao Matos, Petr Cejka

## Abstract

During prophase of the first meiotic division, cells deliberately break their DNA. These DNA breaks are repaired by homologous recombination, which facilitates proper chromosome segregation and enables reciprocal exchange of DNA segments between homologous chromosomes, thus promoting genetic diversity in the progeny^1^. A successful completion of meiotic recombination requires nucleolytic processing of recombination intermediates. Genetic and cellular data implicated a pathway dependent on the putative MLH1-MLH3 (MutLγ) nuclease in generating crossovers, but mechanisms that lead to its activation were unclear^2–4^. Here, we have biochemically reconstituted key elements of this pro-crossover pathway. First, we show that human MSH4-MSH5 (MutSγ), which was known to support crossing over^5–7^, binds branched recombination intermediates and physically associates with MutLγ. This helps stabilize the ensemble at joint molecule structures and adjacent dsDNA. Second, we show that MutSγ directly stimulates DNA cleavage by the MutLγ endonuclease, which demonstrates a novel and unexpected function for MutSγ in triggering crossing-over. Third, we find that MutLγ activity is further stimulated by EXO1, but only when MutSγ is present. Fourth, we also identify the replication factor C (RFC) and the proliferating cell nuclear antigen (PCNA) as additional components of the nuclease ensemble, and show that *S. cerevisiae* strains expressing PIP box-mutated MutLγ present striking defects in forming crossovers. Finally, we show that the MutLγ-MutSγ-EXO1-RFC-PCNA nuclease ensemble preferentially cleaves DNA with Holliday junctions, but shows no canonical resolvase activity. Instead, the multilayered nuclease ensemble likely processes meiotic recombination intermediates by nicking dsDNA adjacent to junction points^8^. Since DNA nicking by MutLγ is dependent on its co-factors, the asymmetric distribution of MutSγ and RFC/PCNA on meiotic recombination intermediates may drive biased DNA cleavage. This unique mode of MutLγ nuclease activation might explain crossover-specific processing of Holliday junctions within the meiotic chromosomal context^3, 9^.

## Introduction

In the prophase of the first meiotic division, SPO11-catalyzed DNA double-strand breaks (DSBs) are repaired by homologous recombination^10^. Joint molecules, such as double Holliday junctions (HJs) or their precursors, arise during recombination and must be ultimately resolved so that chromosomes are properly segregated^1, 11^. A key factor that was implicated in joint molecule metabolism in meiotic cells of most organisms is MutLγ (MLH1-MLH3)^2, 3, 12–15^. Mice lacking MLH1 or MLH3 are infertile^15, 16^, and defects in this pathway may explain infertility in humans^17^. In yeast, MutLγ, together with a group of proteins including MutSγ (Msh4-Msh5) and other meiosis-specific ZMM proteins (for Zip, Msh, Mer), as well as Exo1, is responsible for the majority of crossovers resulting from biased resolution of meiotic recombination intermediates^1, 3, 5^. The meiotic recombination function of yeast MutLγ is dependent on the integrity of the metal binding DQHA(X)_2_E(X)_4_E motif within Mlh3, implicating the nuclease of Mlh3 in resolving recombination intermediates^2, 3, 18, 19, 56^. Despite wealth of genetic and cellular data, the mechanisms that control the MutLγ nuclease and lead to biased joint molecule processing remained undefined.

## Results

### Human MutLγ is an ATP-stimulated endonuclease

To study human MutLγ (hMLH1-hMLH3), we expressed and purified the heterodimer from insect cells (Fig. 1a and Extended Data Fig. 1a,b). Similarly to the mismatch repair (MMR)-specific hMutLα (hMLH1-hPMS2)^20^, the hMLH1-hMLH3 complex non-specifically nicked double-stranded supercoiled DNA (scDNA) in the presence of manganese without any other protein co-factor (Fig. 1b,c, Extended Data Fig. 1c), while almost no activity was observed with magnesium (Extended Data Fig. 1d), which is believed to be the specific metal co-factor^20^. Mutations in the conserved metal binding motif of hMLH3 abolished the endonuclease, indicating that the DNA cleavage activity was intrinsic to the hMutLγ heterodimer (Fig. 1d, see also Extended Data Fig. 1e). ATP promoted the nuclease activity >2-fold (Fig. 1d,e, Extended Data Fig. 1f-h). Experiments with various ATP analogs revealed that ATP hydrolysis by hMLH1-hMLH3 was required for the maximal stimulation of DNA cleavage (Fig. 1f, Extended Data Fig. 1h). The N-termini of both hMLH1 and hMLH3 proteins contain conserved Walker motifs implicated in ATP binding and hydrolysis^21^. To define whether the ATPase of hMLH1, hMLH3 or both subunits of the heterodimer promotes its nucleolytic activity, we prepared the respective hMutLγ variants with mutations in the conserved motifs of either subunit individually or combined (Fig. 1g, Extended Data Fig. 1i)^21^. We observed that the integrity of the ATPase domain of hMLH1, and to a much lesser degree of hMLH3, promoted the nuclease activity of hMLH1-hMLH3 (Fig. 1g, Extended Data Fig. 1j). The hMutLγ complex did not cleave oligonucleotide-based HJ DNA (Extended Data Fig. 1k). Yeast and human MutLγ complexes bind DNA with a preference towards branched structures such as Holliday junctions^18, 22^. The stimulation of DNA cleavage by hMutLγ with ATP can be in part explained by an increased affinity of the heterodimer to DNA when ATP was present (Extended Data Fig. 2a-c). Without ATP, the ATPase-deficient variants of hMLH1-hMLH3 bound DNA indistinguishably from the wild type complex (Extended Data Fig. 2a,d). Our results thus establish that hMLH1-hMLH3 is a nuclease that nicks dsDNA. The endonuclease requires the metal-binding motif within hMLH3 and is promoted upon ATP hydrolysis.

**Figure 1.**
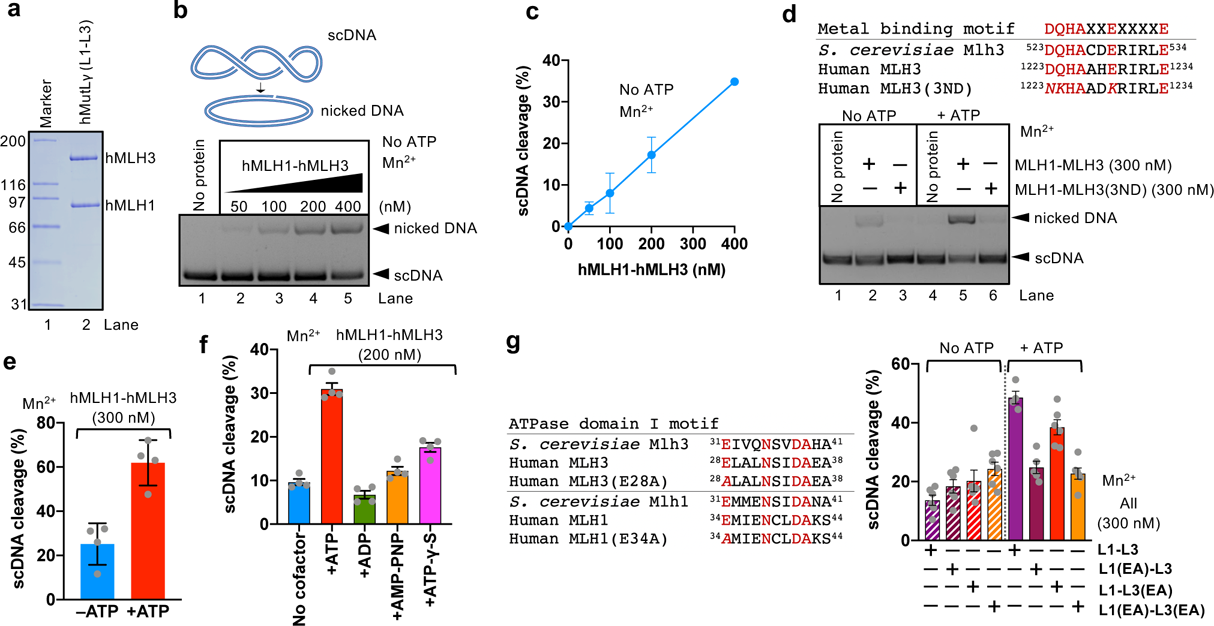
Human hMLH1-hMLH3 is an endonuclease. **a**, Recombinant hMLH1-hMLH3 (L1-L3) used in this study. **b**, Nuclease assay with hMLH1-hMLH3 and pUC19-based negatively supercoiled DNA (scDNA) as a substrate. The reaction with 5 mM manganese acetate was incubated for 60 min at 37 °C. **c**, Quantitation of assays such as in b. Averages shown; error bars, SEM; n=3. **d**, Top, alignment of metal binding motif in yeast and human MLH3. Alanine substitution mutations used in this study are in italics. Bottom, nuclease assay as in b, but with wild type hMLH1-hMLH3 (L1-L3) or nuclease-dead L1-L3(3ND) (mutations D1223N, Q1224K, E1229K) hMLH1-hMLH3 variants, without or with ATP (0.5 mM). **e**, Quantitation of nuclease assays with hMLH1-hMLH3 without or with ATP (0.5 mM), in the presence of manganese (5 mM). Averages shown; error bars, SEM; n=4. **f**, Quantitation of assays as in Extended Data Fig. 1h, supplemented with various nucleotide co-factors and their analogs (0.5 mM). Averages shown; error bars, SEM; n=4. **g**, Left, alignment of MLH1 and MLH3 ATPase domains from humans and yeast. Conserved residues are highlighted in red. Alanine substitutions in MLH3 and MLH1 used in this study are in italics. Right, quantitation of assays as in Extended Data Fig. 1j, without or with ATP (0.5 mM), with either wild type hMLH1-hMLH3, L1-L3; hMLH1(E34A)-hMLH3, L1(EA)-L3; hMLH1-hMLH3(E28A), L1-L3(EA), or hMLH1(E34A)-hMLH3(E28A), L1(EA)-L3(EA). Averages shown; error bars, SEM; n=3.

**Figure 2.**
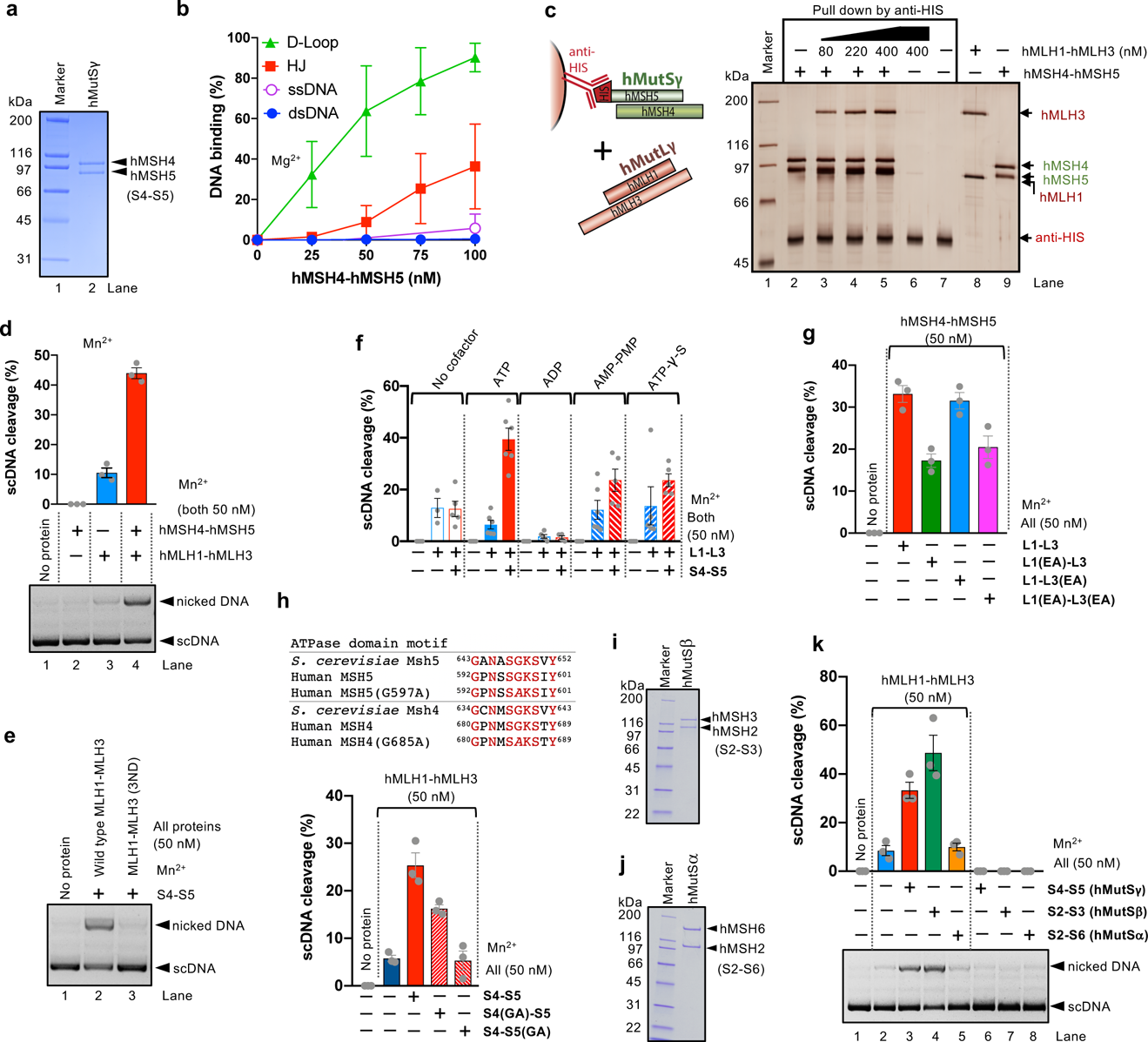
hMSH4-hMSH5 directly promotes the endonuclease activity of hMLH1-hMLH3. **a**, Recombinant hMSH4-hMSH5 (S4-S5) used in this study. **b**, Quantitation of DNA binding assays such as shown in Extended Data Fig. 3a. Averages shown; error bars, SEM; n=3. **c**, Protein interaction assays with immobilized hMSH4-hMSH5 (bait) and hMLH1-hMLH3 (prey). Lanes 8 and 9 show recombinant proteins loaded as controls. The 10% polyacrylamide gel was stained with silver. **d**, Nuclease assays with pFR-Rfa2 5.6 kbp-long scDNA and hMLH1-hMLH3 and hMSH4-hMSH5, as indicated. The assays were carried out at 30 °C in the presence of 5 mM manganese acetate and 0.5 mM ATP. A representative experiment is shown at the bottom, a quantitation (averages shown; n=3; error bars, SEM) at the top. **e**, Nuclease assays with hMSH4-hMSH5 (S4-S5) and either wild type hMLH1-hMLH3 (L1-L3) or nuclease-dead hMLH1-hMLH3 (D1223N, Q1224K, E1229K) (L1-L3[3ND]). The assays were carried out at 30 °C in the pres-ence of 5 mM manganese acetate and 0.5 mM ATP. **f**, Quantitation of nuclease assays with hMLH1-hMLH3 (L1-L3) and hMSH4-hMSH5 (S4-S5), as indicated, in the presence of various nucleotide co-factors or their analogs (2 mM). The assays were carried out at 30 °C in the presence of 5 mM manganese acetate. Averages shown; error bars, SEM; n≥4. **g**, Quantitation of nuclease assays as shown in Extended Data Fig. 5d, with variants of hMLH1-hMLH3 (L1-L3), deficient in ATP hydrolysis, without or with hMSH4-hMSH5 (S4-S5). See also Fig. 1g. Averages shown; error bars, SEM; n=3. The assays were carried out at 37 °C in the presence of 5 mM manganese acetate and 0.5 mM ATP. **h**, Top, alignment of MSH5 and MSH4 ATPase domains from humans and yeast. Conserved residues are highlighted in red. Alanine substitutions in MSH5 and MSH4 used in this study are in italics. Bottom, quantitation of nuclease assays as shown in Extended Data Fig. 5e, with variants of hMSH4-hMSH5 (S4-S5), deficient in ATP hydrolysis, and hMLH1-hMLH3. Averages shown; error bars, SEM; n=3. The assays were carried out at 30°C in the presence of 5 mM manganese acetate and 0.5 mM ATP. **i**, Recombinant hMutS*β* (hMSH2-hMSH3) used in this study. **j**, Recombinant hMutS*α* (hMSH2-hMSH6) used in this study. **k**, Nuclease assays with hMLH1-hMLH3, hMSH4-hMSH5 (S4-S5), and hMSH2-hMSH3 (S2-S3) or hMSH2-hMSH6 (S2-S6), as indicated. The assays were carried out at 30 °C in the presence of 5 mM manganese acetate and 0.5 mM ATP. A representative experiment is shown at the bottom, a quantitation (averages shown; n=3; error bars, SEM) at the top.

### MutLγ and MutSγ interact and stabilize each other at DNA junctions

Previously, recombinant hMSH4-hMSH5 was shown to bind HJs^6^. We found that the human and yeast MutSγ complexes bound even better precursors of HJs such as D-loops (Fig. 2a-b, Extended Data Fig. 3a-d). This is in agreement with a proposed early function of MutSγ and other ZMM proteins to stabilize nascent strand invasion intermediates that mature into single-end invasions, which helps ensure their crossover designation^7, 23^. In contrast, single-stranded DNA (ssDNA) or dsDNA was not bound by MutSγ, establishing thus the binding preference of the heterodimer to branched DNA structures (Fig. 2b, Extended Data Fig. 3a-d), similarly to MutLγ^18, 22^. Electrophoretic mobility shift assays demonstrated that the MutSγ and MutLγ complexes moderately stabilized each other at DNA junctions, which required the interplay of the cognate heterodimers (Extended Data Fig. 4a-d). Accordingly, the respective human or yeast MutSγ and MutLγ complexes directly physically interact (Fig. 2c and Extended Data Fig. 4e-g)^24^. The very slow migration of the protein-DNA complexes was indicative of multiple units of the heterocomplexes bound to the DNA substrate (Extended Data Fig. 4h,i), as shown previously for yeast MutLγ^18^. We note that the presence of DNA junctions was essential for stable DNA binding (Extended Data Fig. 4a,c), which supports a model where a branched DNA structure serves as a nucleation point for a hMutSγ-hMutLγ filament that then extends to the adjacent dsDNA arms^25^.

**Figure 3.**
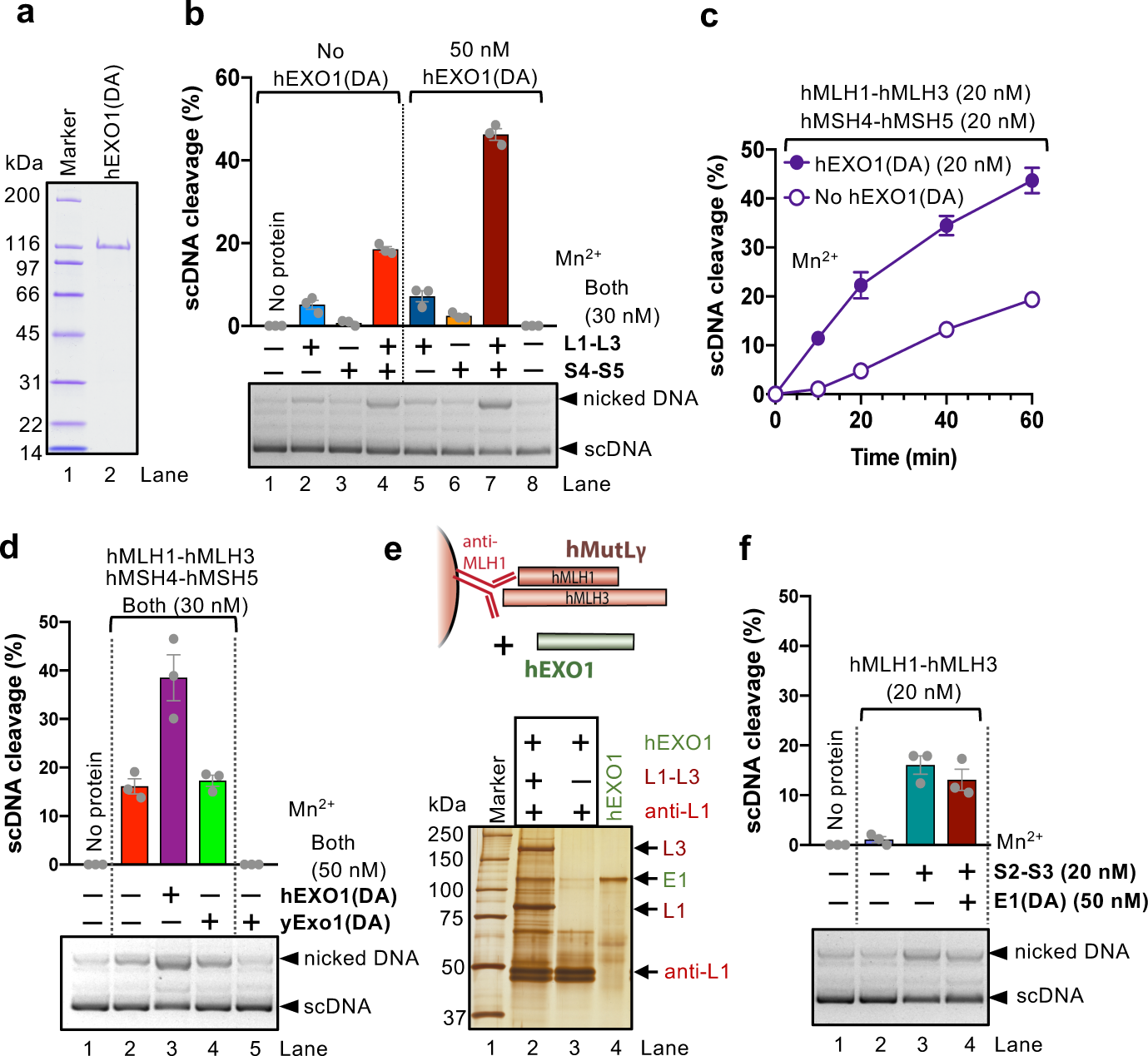
hEXO1 promotes the nuclease activity of hMLH1-hMLH3 when in complex with hMSH4-hMSH5. **a**, Recombinant hEXO1(D173A) used in this study. **b**, Nuclease assays with hMLH1-hMLH3 (L1-L3) and hMSH4-hMSH5 (S4-S5), as indicated, without (left) or with hEXO1(DA) (right). The assays were carried out at 30 °C in the presence of 5 mM manganese acetate and 0.5 mM ATP. A representative experiment is shown at the bottom, a quantitation (averages shown; n=3; error bars, SEM) at the top. **c**, Quantitation of kinetic nuclease assays with hMLH1-hMLH3 and hMSH4-hMSH5, without or with hEXO1(DA). The assays were carried out at 30°C in the presence of 5 mM manganese acetate and 2 mM ATP. Averages shown; error bars, SEM; n=3. **d**, Nuclease assays with hMLH1-hMLH3, hMSH4-hMSH5 with either human hEXO1(D173A) or yeast yExo1(D173A), as indicated. The assays were carried out at 30 °C in the presence of 5 mM manganese acetate and 0.5 mM ATP. A representative experiment is shown at the bottom, a quantitation (averages shown; n=3; error bars, SEM) at the top. **e**, Protein interaction assays with immobilized hMLH1-hMLH3 (L1-L3, bait) and hEXO1 (E1, prey). Lane 3, recombinant hEXO1 was loaded as a control. The 10% polyacrylamide gel was stained with silver. **f**, Nuclease assays with hMLH1-hMLH3, hMSH2-hMSH3 (S2-S3) and hEXO1(DA), as indicated. The assays were carried out at 30 °C in the presence of 5 mM manganese acetate and 0.5 mM ATP. A representative experiment is shown at the bottom, a quantitation (averages shown; n=3; error bars, SEM) at the top.

**Figure 4.**
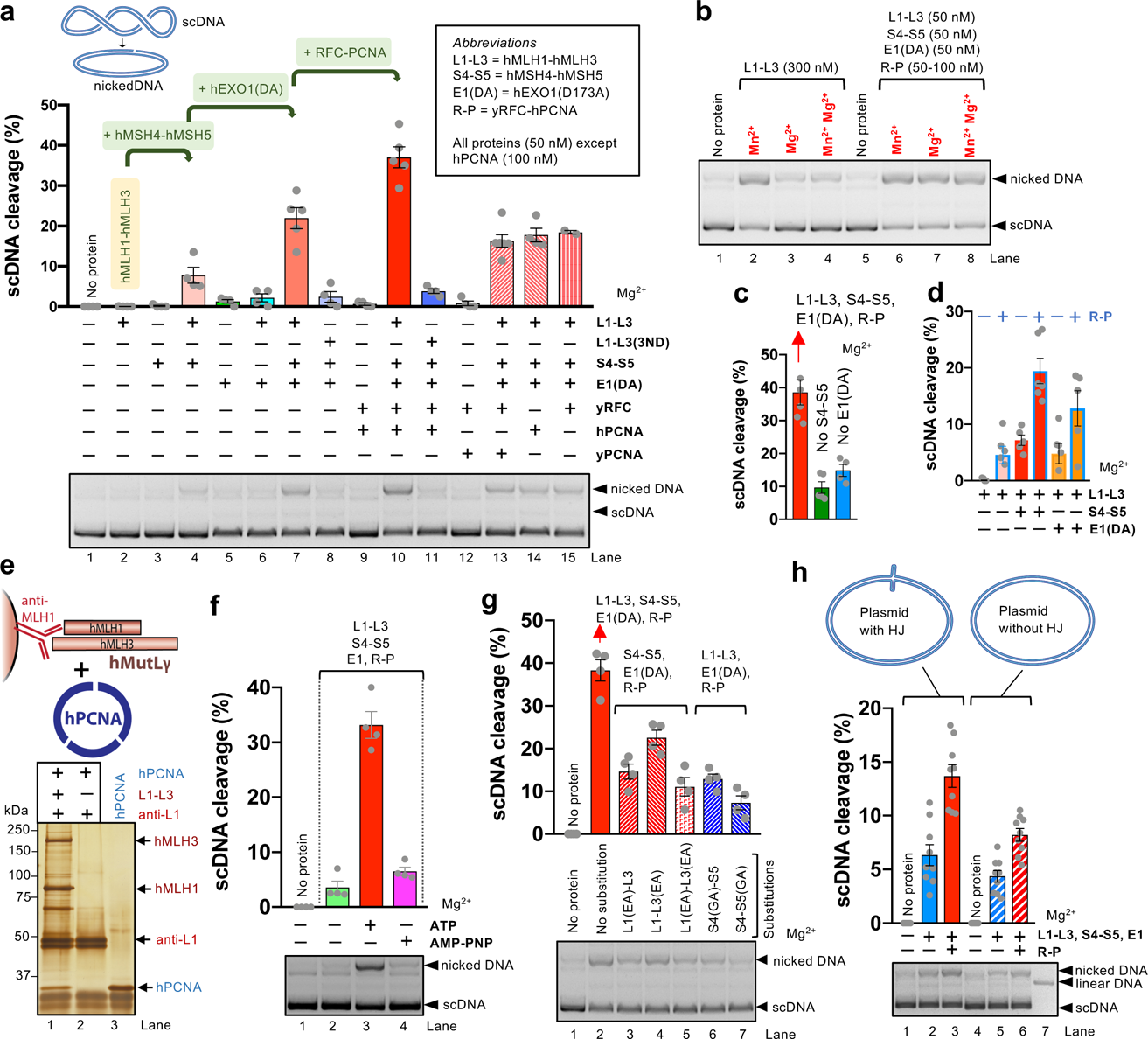
RFC-PCNA promote DNA cleavage by the hMutLγ-hMutSγ-hEXO1(D173A) ensemble. **a**, Nuclease assays with scDNA and indicated proteins (all 50 nM, except hPCNA, 100 nM) was carried out with 5 mM magnesium acetate and 2 mM ATP at 37 °C. A representative experiment is shown at the bottom, a quantitation (averages shown; n≥4; error bars, SEM) at the top. **b**, Representative nuclease assays carried out with 5 mM magnesium and/or manganese acetate, as indicated, with indicated recombinant proteins, containing 2 mM ATP and incubated at 37 °C. **c**, Nuclease reactions containing hMLH1-hMLH3 (L1-L3, 50 nM); hMSH4-hMSH5 (S4-S5, 50 nM); hEXO1(D173A) (E1(DA), 50 nM) and yRFC-hPCNA (R-P, 50-100 nM, respectively) (column 1), without hMSH4-hMSH5 (column 2) or without hEXO1(D173A) (column 3). Reactions were carried out with 5 mM magnesium acetate and 2 mM ATP at 37 °C. Averages shown; error bars, SEM; n≥4. **d**, Nuclease reactions with hMLH1-hMLH3 (L1-L3, 50 nM); hMSH4-hMSH5 (S4-S5, 50 nM); hEXO1(D173A) (E1(DA), 50 nM) and yRFC-hPCNA (R-P, 50-100 nM, respectively), as indicated. Reactions were carried out with 5 mM magnesium acetate and 2 mM ATP at 37 °C. Averages shown; error bars, SEM; n≥5. **e**, Protein interaction assays with immobilized hMLH1-hMLH3 (bait) and hPCNA (prey). Lane 3, recombinant hPCNA was loaded as a control. The 10% polyacrylamide gel was stained with silver. **f**, Nuclease reactions as in panel a, lane 10, but either without ATP, with ATP or with AMP-PNP (2 mM). Averages shown; error bars, SEM; n=4. **g**, Nuclease reactions with hMLH1-hMLH3 (L1-L3, 50 nM); hMSH4-hMSH5 (S4-S5, 50 nM); hEXO1(D173A), (E1(DA), 50 nM) and yRFC-hPCNA (R-P, 50-100 nM, respectively), lane 2. Lanes 3-7 contain instead hMLH1-hMLH3 or hMSH4-hMSH5 variants deficient in ATP hydrolysis, as indicated. See Extended Data Fig. 5d,e for specific mutations. Reactions were carried out with 5 mM magnesium acetate and 2 mM ATP at 37 °C. Averages shown; error bars, SEM; n=4. **h**, Representative nuclease reactions as in panel a, but with 3.5 kbp-long dsDNA either containing (left) or not (right) DNA repeat forming HJ-like cruciform DNA. Averages shown; error bars, SEM; n=9.

### hMutSγ directly promotes the endonuclease activity of hMutLγ

Previous *in vivo* experiments implicated MutSγ in the stabilization of nascent DNA joint molecules early in the meiotic pro-crossover pathway^6, 24, 26^, but whether MutSγ is directly involved later in nucleolytic processing was not clear. Using our reconstituted system, we observed ∼3-fold stimulation of hMLH1-hMLH3 endonuclease by hMSH4-hMSH5 (Fig. 2d, Extended Data Fig. 5a-c), which was dependent on the hMLH3 metal binding motif (Fig. 2e). ATP promoted DNA cleavage by the hMutSγ-hMutLγ ensemble, and as in reactions with hMutLγ alone, maximal nuclease activity was observed when ATP hydrolysis was possible (Fig. 2f). The ATP binding/hydrolysis motifs in hMLH1 and hMSH5 were both crucial, while the motif in hMSH4 was less important and in hMLH3 appeared dispensable (Fig. 2g,h, Extended Data Fig. 5d,e). The ATPase motif mutations in hMSH4 or hMSH5 instead did not affect the capacity of the two subunits to form a complex or bind DNA (Extended Data Fig. 5f,g). The stimulatory effect was likely dependent on direct physical interactions between the cognate heterodimers, as yeast Msh4-Msh5 did not promote the nuclease of human MutLγ (Extended Data Fig. 5h). The hMutSγ-hMutLγ complex cleaved similarly both supercoiled and relaxed DNA (Extended Data Fig. 5i) and exhibited no detectable structure-specific nuclease or resolvase activity (Extended Data Fig. 5j,k).

**Figure 5.**
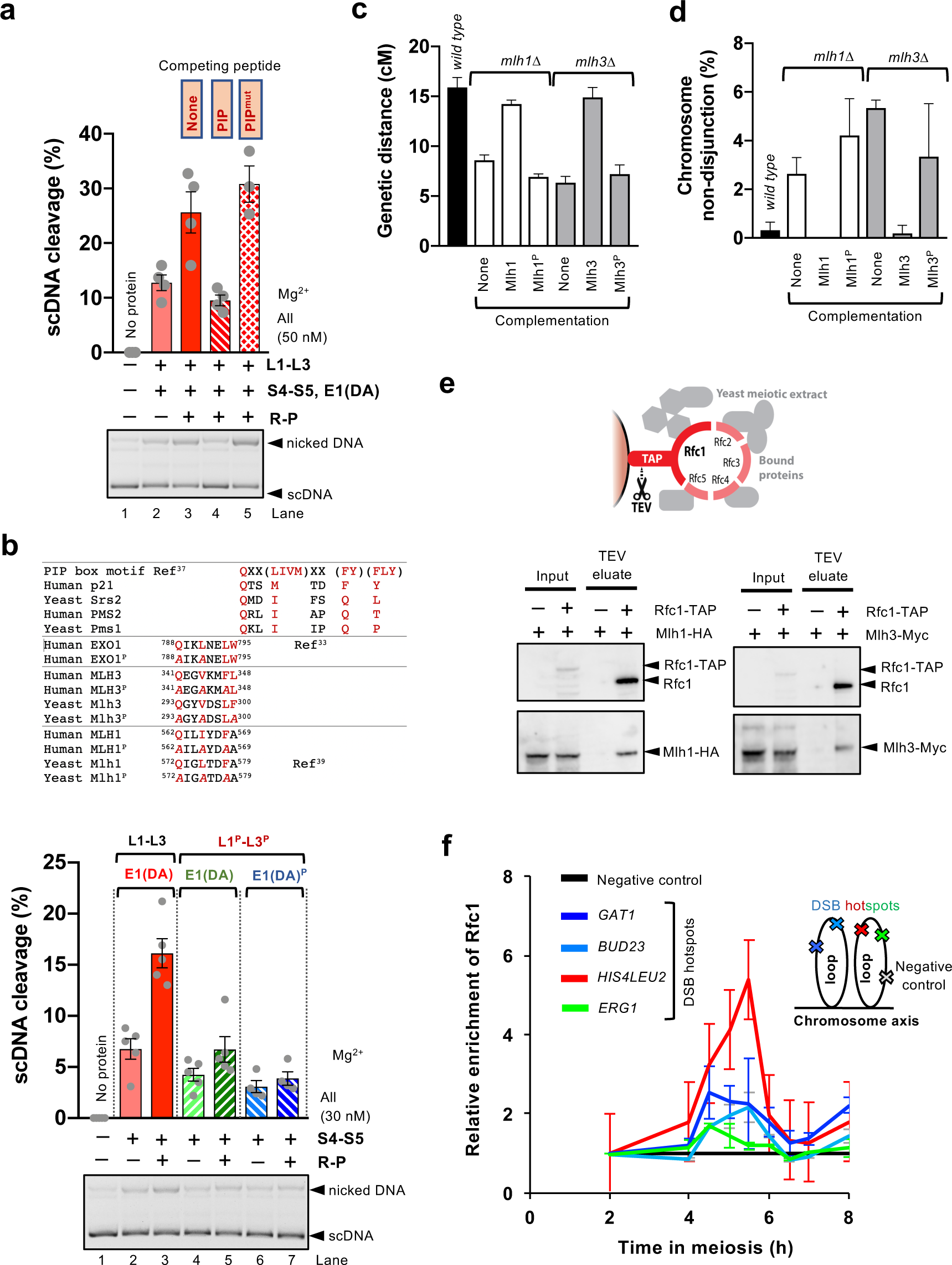
The stimulation of MLH3 nuclease ensemble requires a PIP box motif and is conserved in evolution. **a**, Nuclease assays with hMLH1-hMLH3, L1-L3 (50 nM); hMSH4-hMSH5, S4-S5 (50 nM); hEXO1(DA), E1(DA) (50 nM) and yRFC-hPCNA, R-P (50-100nM), as indicated, with 5 mM magnesium acetate and 2 mM ATP at 37 °C. The reactions were supplemented with a p21^22^ PIP-box wild type or mutated control peptide, where indicated (670 nM, ∼5-fold over *K_d_* of wild type peptide for PCNA). Averages shown; error bars, SEM; n=4. **b**, Top, alignment of PIP-box like motifs from various human or yeast (*S. cerevisiae*) proteins. Residues more likely to be conserved are highlighted in red. Wild type human and yeast EXO1, MLH3/Mlh3 and MLH1/Mlh1 were mutated to create respective (^P^) variants with indicated residue substitutions (*A*). Bottom, nuclease assays with hMLH1-hMLH3, L1-L3 (50 nM); hMSH4-hMSH5, S4-S5 (50 nM); hEXO1(DA), E1(DA) (50 nM) and yRFC-hPCNA, R-P (50-100nM), as indicated, with 5 mM magnesium acetate and 2 mM ATP at 37 °C. Where indicated, wild type hMLH1 was replaced with MLH1^P^ (Q562A, I565A, F568A), wild type hMLH3 with hMLH3^P^ (Q341A, V344A, F347A), and E1(DA), hEXO1(D173A), with E1(DA)^P^, hEXO1 (D173A, Q788A, L791A). Averages shown; error bars, SEM, n=5. **c**, Recombination frequency, expressed as a map distance in centimorgans, was assayed in the wild type strain, *mlh1Δ* and *mlh3Δ*, and in strains complemented with a construct expressing wild type Mlh1, Mlh1^P^ (Q572A-L575A-F578A) or Mlh3^P^ (Q293A-V296A-F300A). Averages shown; error bars, SD; n≥900 from 3 biological replicates for each genotype. **d**, Frequency of chromosome VIII non-disjunction in strains as described in panel c. Averages shown; error bars, SD; n≥900 from 3 biological replicates for each genotype. **e**, A pulldown of TAP-tagged yRfc1-5 and associated proteins from meiotic cell extracts from p*CUP1*-*IME1* cells 5 h 30 min after the induction of meiosis. The presence of Mlh1-HA and Mlh3-Myc in the TEV eluate was analyzed by Western blotting. **f**, Rfc1-TAP levels at the indicated meiotic DSB hotspots relative to a negative control site (*NFT1)* were assessed by ChIP and qPCR during a meiotic time-course (synchronized *pCUP1-IME1* cells). Averages shown; error bars, SD; n=2. The cartoon illustrates the position of sites analyzed by qPCR relative to the meiotic chromosome structure.

To determine whether the MMR-specific human MutS homologue complexes could also promote hMutLγ, we supplemented hMLH1-hMLH3 reactions with recombinant hMutS*α* (hMSH2-hMSH6) or hMutS*β* (hMSH2-hMSH3) (Fig. 2i-j). hMutS*β*, but not hMutS*α*, could also stimulate the hMLH1-hMLH3 nuclease (Fig. 2k). This agrees with previous experiments showing that yeast MutLγ could partially substitute MutLα in the repair of insertion/deletion mismatches in MMR^27^. These data also underpin the involvement of hMutLγ in the metabolism of trinucleotide repeats linked to several neurodegenerative diseases, as well as rare hMLH3 mutations found in patients with hereditary nonpolyposis colorectal cancer (HNPCC)/Lynch syndrome characterized by microsatellite instability^28–32, 56^.

### hEXO1 promotes the nuclease activity of hMutSγ-hMutLγ

Genetic experiments with budding yeast revealed a structural (nuclease-independent) function of Exo1 in the Mlh1-Mlh3 pro-crossover pathway^4^. The effect was dependent on its direct interaction with the Mlh1 subunit of the yMutLγ heterodimer^4, 33^, but it was unclear whether the interplay directly affects the Mlh3 endonuclease, and whether this function is conserved in higher eukaryotes. While hEXO1 is likely one of the nucleases that function in MMR to exonucleolytically remove the DNA stretch containing the mismatch, its role in the initial endonucleolytic cleavage catalyzed by hMutLα was not reported^20^. To test the effect of hEXO1 on the nuclease of hMLH1-hMLH3, we used the nuclease-deficient hEXO1(DA) variant to prevent degradation of the resulting nicked DNA (Fig. 3a, Extended Data Fig. 6a). We observed no effect of hEXO1(DA) on the nuclease of hMLH1-hMLH3 alone, but hEXO1(DA) promoted DNA cleavage ∼2-3-fold when hMSH4-hMSH5 was present (Fig. 3b,c and Extended Data Fig. 6b). More than 40% DNA cleavage was observed using 20 nM concentration of the multi-protein ensemble (Fig. 3c). Compared to DNA cleavage efficiency by 400 nM hMLH1-MLH3 alone in reactions without ATP (Fig. 1b,c), this corresponds to >20-fold stimulation of nuclease activity by the respective co-factors.

In contrast to hMSH4-hMSH5 that stabilized hMLH1-hMLH3 on DNA, we detected no such capacity of hEXO1(DA) (Extended Data Fig. 6c,d). Yeast Exo1(DA) could not substitute human EXO1(DA) in the nuclease assays (Fig. 3d), in agreement with a direct physical interaction between human hEXO1(DA) and hMLH1-hMLH3 (Fig 3e). We also note that the ensemble was inefficient in cleaving DNA opposite to nicks, showing that nicks are unlikely to direct the endonuclease (Extended Data Fig. 6e). Finally, hEXO1(DA) did not promote the nuclease of hMLH1-hMLH3 in conjunction with the MMR-specific hMSH2-hMSH3 complex (Fig. 3f), indicating that hEXO1 likely specifically promotes the endonuclease activity of hMutSγ-hMutLγ involved in meiotic recombination.

### RFC-PCNA promote the activity of the MutSγ-hEXO1(DA)-MutLγ nuclease ensemble

We next set out to test whether hMLH1-hMLH3 with its co-factors can catalyze DNA cleavage under physiological conditions in magnesium. While almost no nuclease activity of hMLH1-hMLH3 alone in magnesium was observed, weak nicking was seen in the presence of hMSH4-hMSH5, and the reactions were further stimulated by hEXO1(DA) (Fig. 4a). As RFC-PCNA are known to promote the hMLH1-hPMS2 (hMutL*α*) endonuclease in MMR^20^, we tested for their effect on hMutLγ. Notably, we observed additional ∼2-fold stimulation of DNA cleavage by the nuclease ensemble when RFC-PCNA complex was present (Fig. 4a, compare lanes 7 and 10, Extended Data Fig. 7a-c). The reactions with hMutLγ-hMutSγ-hEXO1-RFC-PCNA were dependent on the integrity of the hMLH3 metal-binding motif (Fig. 4a, lane 11). Yeast RFC is capable of loading human PCNA, and could readily substitute human RFC in reconstituted MMR reactions^20^. In accord, we observed the stimulatory effect on the hMLH1-hMLH3 ensemble when using yeast RFC and human PCNA, but not when using yeast RFC and yeast PCNA (Fig. 4a, lane 13). Also, no stimulation was detected when RFC was omitted from the reaction mixture containing human PCNA (Fig. 4a, lane 14), indicating that PCNA must likely be actively loaded onto DNA by RFC, as during MMR^20, 34, 35^. Accordingly, PCNA is known to be efficiently loaded onto intact negatively supercoiled DNA^34^. In summary, we show that while hMutLγ *per se* is a poor nuclease that requires manganese, hMutSγ, hEXO1 and RFC-PCNA activate it to cleave efficiently in a buffer containing physiological magnesium, and the reaction is no longer stimulated by adding manganese (Fig. 4b, see also Extended Data Fig. 7d). The omission of hMSH4-hMSH5 or hEXO1(DA) resulted in a strong reduction of the ensemble nuclease activity (Fig. 4c), demonstrating the requirement for the multiple co-factors to simultaneously stimulate hMutLγ. RFC-PCNA could also promote the nuclease of hMLH1-hMLH3 alone, although to a much lesser extent (Fig. 4d), suggesting that hMutSγ and hEXO1(DA) are not strictly required to mediate the stimulatory effect of RFC-PCNA. In accord, we found that hMutLγ directly physically interacts with hPCNA (Fig. 4e).

ATP was necessary for the nuclease activity of the ensemble, and could not be replaced by ADP or AMP-PNP, showing that ATP hydrolysis was required (Fig. 4f, Extended Data Fig. 7e). In contrast to the reactions in manganese, the integrity of the ATPase motifs of all four hMutSγ and hMutLγ subunits was required for maximal cleavage activity (Fig. 4g), in agreement with meiotic defects of the corresponding ATPase-deficient yeast mutant strains^21, 23^.

Notably, the nuclease ensemble preferentially cleaved plasmid-length DNA with palindromic repeats forming a HJ-like structure (Fig. 4h), in agreement with the binding preference of the hMutSγ and hMutLγ heterodimers to these recombination intermediates. However, the activity of the complex on the cruciform DNA primarily yielded nicked products (Fig. 4h), unlike canonical HJ resolvases that give rise to linear DNA upon concerted cleavage of both DNA strands at the junction points^36^. We note that we did not observe any cleavage of model HJ or D-loop oligonucleotide-based substrates (Extended Data Fig. 7f). Our data suggest that the hMutLγ ensemble processes recombination intermediates by resolution-independent nicking, in agreement with results obtained from sequencing of heteroduplex DNA arising during meiosis in yeast cells, which indicated cleavage by nicking some distance away from the DNA junction points^8^.

Interactions with PCNA are often mediated by a PCNA-interacting peptide (PIP) motif^37^. We supplemented the hMutLγ ensemble nuclease assays with a PIP-box peptide derived from p21^38^, or a control peptide with key residues mutated. The competing PIP-box peptide eliminated the stimulatory effect of RFC-PCNA, while the control peptide had no effect (Fig. 5a, concentration of the peptide was ∼5-fold over *K_d_* for PCNA^38^), demonstrating that the PCNA function in stimulating the nuclease of the hMutLγ ensemble is dependent on an interaction via a PIP-box like motif. We next analyzed several mutants of conserved PIP-box-like sequences in MLH1, MLH3 and EXO1(DA)^35, 39, 40^ (Extended Data Fig. 5b, top). The respective mutations did not notably affect the nuclease reactions *per se* without PCNA or the capacity to bind DNA (Extended Data Fig. 8a-d), but the mutants became partly refractory to stimulation by RFC-PCNA, in particular when the PIP-box-like mutations of multiple factors were combined (Fig. 5b, Extended Data Fig. 8c,e). Furthermore, the corresponding mutations in the yeast homologues of Mlh1 and Mlh3 (see scheme at the top of Fig. 5b) resulted in meiotic defects, as indicated by a decrease in the frequency of crossovers at *CEN8*-*THR1* leading to chromosome non-disjunction and reduced spore viability (Fig. 5c,d, Extended Data Fig. 9a). We also found yeast Mlh1 and Mlh3 as a part of a complex with yRfc1 in meiotic cells at the time of joint molecule resolution (Fig. 5e). Finally, using chromatin immunoprecipitation and synchronous meiotic yeast cultures, we observed an enrichment of yRfc1 at both natural and engineered DSB hotspots in late meiosis at the time when joint molecules are resolved into crossovers (Fig. 5f), which coincides with the accumulation of Mlh3 at the same hotspots (V. Borde, manuscript in preparation). The accumulation of yRfc1 at sites of recombination was independent of Mlh3, suggesting it may be present from an earlier step of DNA synthesis (Extended Data Fig. 9b). The function of RFC-PCNA in promoting meiotic crossovers in the MLH3 pathway is thus likely conserved in evolution.

## Discussion

The MutLγ nuclease in most organisms functions in a pathway that processes a subset of meiotic joint DNA molecules into crossover recombination products, and was thus proposed to represent a crossover-specific resolvase^3^. Here we show that hMSH4-hMSH5, hEXO1 and RFC-PCNA strongly promote the MutLγ (hMLH1-hMLH3) nuclease, but we failed to detect any canonical HJ resolvase activity. Rather, our data suggest that the nuclease ensemble processes joint molecule intermediates by biased resolution-independent nicking of dsDNA in the vicinity of HJs. Since HJs are symmetric and their resolution can yield both crossovers and non-crossovers^41^, how is the crossover bias established? As hMSH4-hMSH5 likely stabilizes asymmetric HJ precursors^5^, it is anticipated that the heterodimer will be ultimately present asymmetrically later at the mature joint molecules containing HJs (Extended Data Fig. 10). Similarly, PCNA may be loaded asymmetrically at joint molecules during DNA synthesis by polymerase *δ*^42^, or at strand discontinuities before the ligation of double HJs takes place. Asymmetric localization of MSH4-MSH5 was indeed directly observed by structured-illumination microscopy in *Caenorhabditis elegans*^43^. Although the crossover-specific resolution of DNA junctions in worms is independent of MLH1-MLH3 and depends on other nucleases, the asymmetric presence of nuclease co-factors might Represent a general mechanism to promote biased joint molecule resolution. In yeast, the Yen1 nuclease that was artificially activated instead of Mlh1-Mlh3 also led to crossover-biased resolution, arguing that the nature of the substrate within the meiotic chromosome context, rather than the Mlh1-Mlh3 nuclease *per se,* directs the biased resolution^9^. We propose that the asymmetric presence of RFC-PCNA, MSH4-MSH5 or additional co-factors at joint molecules might provide the signal to guarantee the biased, crossover-specific processing by the MutLγ nuclease (Extended Data Fig. 10), and the mechanism can be broadly applicable to other organisms that do not possess MutLγ.

## Acknowledgements

This work was supported by grants from the Swiss National Science Foundation (31003A_17544) and European Research Council (681-630) to P.C., Institut Curie and CNRS to V.B., by Agence Nationale de la Recherche (ANR-15-CE11-0011) to V.B. and J.-B.C. We thank Josef Jiricny (ETH Zurich) and members of the Cejka laboratory for helpful comments on the manuscript and Neil Hunter for communicating results prior to publication.

## Author contributions

E.C., A.S., R.A. and P.C. planned, performed and analyzed the majority of the experiments and wrote the paper. L. R. and A. A. performed most of the experiments with yeast recombinant proteins and electrophoretic mobility shift assays. N. W. performed experiments to define simultaneous DNA binding by MLH1-MLH3 and MSH4-MSH5. J. H. performed experiments with yeast *mlh1* and *mlh3* variants mutated in PIP-box-like sequences, and the data were analyzed together with J. M. Chip experiments and Rfc1-Mlh1 and Rfc1-Mlh3 pulldown assays were carried out by C. A., the data were analyzed together with V. B. J-B.C. helped prepare the MLH1-MLH3 expression construct and designed experiments with the PIP-box peptide. X. A-G. and E.R.H. prepared the MSH4-MSH5 expression construct. All authors contributed to prepare the final version of the manuscript.

## Methods

### Preparation of expression vectors

To prepare the hMSH4-hMSH5 expression vector, the hMSH4-STREP and hMSH5-8xHIS constructs were codon-optimized for expression in *Spodoptera frugiperda Sf*9 cells and synthesized (GenScript). The genes were amplified by PCR using M13 forward and reverse primers (see Extended Data Table 1 for sequences of all oligonucleotides) and digested with *Sal*I and *Hin*dIII (for hMSH4) or *Sma*I and *Kpn*I (for hMSH5) restriction endonucleases (New England Biolabs). Digested fragments were ligated into corresponding sites in pFBDM (Addgene) to obtain pFBDM-hMSH4co-STREP and pFBDM-hMSH5co-HIS, respectively. Both plasmids were then digested with *Bam*HI and *Hin*dIII (New England Biolabs) and ligated to generate pFBDM-hMSH4co-STREP-hMSH5co-HIS. To prepare expression constructs coding for the ATPase variants of hMSH4 and hMSH5, the respective conserved residues in the Walker A motifs (see ref^23^) were mutated by QuikChange II site-directed mutagenesis kit (Agilent Technologies). To prepare MSH4G685A, the pFB-hMSH4co-STREP-hMSH5co-HIS vector was mutated with primers HMSH4G685A_FO and HMSH4G685A_RE. This created pFB-hMSH4coG685A-STREP-hMSH5co-HIS. To prepare MSH5G597A, HMSH5G597A_FO and HMSH5G597A_RE primers were used to create pFB-hMSH4co-STREP-hMSH5coG597A-HIS. We also prepared a construct combining both mutations, but the resulting mutant complex was not stable and could not be purified.

To prepare the hMLH1 and hMLH3 expression vectors, both genes were amplified by PCR from pFL-his-MLH3co-MLH1co containing both *hMLH1* and *hMLH3* genes, which were codon-optimized for insect cell expression. To amplify *hMLH1*, FLAG-hMLH1co_FO and hMLH1co_RE primers were used. The PCR product was digested with *Nhe*I and *Xba*I (New England Biolabs) and inserted in pFB-MBP-MLH3-his^18^ creating pFB-FLAG-hMLH1co (the sequence of *hMLH3* was removed during this step). Similarly, hMLH3 was amplified using MLH3co_FO and MLH3co_RE. The PCR product was digested with *Nhe*I and *Xma*I (New England Biolabs) and inserted into pFB-MBP-MLH3-HIS, generating pFB-MBP-hMLH3co. The sequence of non-optimized *hMLH3* was removed during this step. The consensus metal-binding motif in hMLH3 is <u>DQHA</u>AD<u>E</u> (conserved residues underlined)^2, 20^. To prepare the nuclease-dead variant, the sequence of wild type *hMLH3* in pFB-MBP-MLH3co-HIS was mutated using primers HMLH33ND_FO and HMLH33ND_RE. This created a sequence with 3 point mutations including D1223N, Q1224K and E1229K (*NK*HAAD*K*, mutated residues in italics), and the resulting vector was pFB-MBP-MLH3co3ND-HIS. We note that a single point mutant hMLH1-hMLH3(D1223N) retained ∼10% activity in reactions with manganese (Extended Data Fig. 1e), and was therefore not further used in this work. To disrupt the ATPase of hMLH1^21^, the pFB-FLAG-hMLH1co was mutated using primers HMLH1E34A_FO and HMLH1E34A_RE. This created pFB-FLAG-MLH1E34A. To mutate the corresponding conserved residue in hMLH3, the pFB-HIS-MBP-MLH3co was mutated using primers HMLH3E28A_FO and HMLH3E28A_RE. This created pFB-HIS-MBP-MLH3E28A. To prepare the hMLH1^P^ variant, the pFB-FLAG-hMLH1co plasmid was mutated using HMLH1_PIP1_3AFO and HMLH1_PIP1_3ARE primers. To prepare the hMLH3^P^ variant, the pFB-FLAG-hMLH3co plasmid was mutated using HMLH3_PIP_3AFO and HMLH3_PIP_3ARE primers.

To prepare pFB-hEXO1-FLAG, the sequence coding for wild type hEXO1 (or hEXO1[DA], containing the D173A mutation inactivating its nuclease) was amplified by PCR using primers HEXO1_FO and HEXO1_RE, and respective vectors (a kind gift from Stefano Ferrari, University of Zurich)^44^ as templates. The PCR products were digested by *Bam*HI and *Xma*I (New England Biolabs), and cloned into corresponding sites in pFB-MBP-Sae2-HIS^45^ (the sequence of MBP-Sae2 was removed during the process, FLAG-tag was added to the C-terminus and a HIS-tag from the original construct was not translated due to a Stop codon). The

To prepare pFB-hMSH2-FLAG, the sequence coding for hMSH2 was amplified from pFB-hMSH2^46^ using primers HMSH2FLAG_FO and HMSH2FLAG_RE. The PCR product was digested by *Bam*HI and *Xho*I (New England Biolabs), and cloned into corresponding sites in pFB-MBP-Sae2 (the sequence of MBP-Sae2 was removed during the process, FLAG-tag was added to the C-terminus of hMSH2 and a HIS-tag from the original construct was not translated due to a Stop codon).

To prepare pFB-HIS-yMLH1, pFB-GST-yMLH1^18^ was digested using *Bam*HI (New England Biolabs) to remove the GST tag. This procedure left behind a single *Bam*HI site. Two complementary oligonucleotides His-For and His-Rev were annealed to each other, and cloned into the *Bam*HI site. This introduced a sequence coding for 8xHIS tag before the yeast *MLH1* gene creating pFB-HIS-yMLH1.

To prepare pFB-MBP-yMLH3, a termination codon was introduced after the yMLH3 gene in pFB-MBP-yMLH3-HIS^18^ so that the HIS-tag would not be translated with the yMlh3 protein. This was carried out by site-directed mutagenesis using forward primer 329 and reverse primer 330.

To prepare the expression vector for yMsh4, the yeast *MSH4* gene was amplified from the genomic DNA of the *S. cerevisiae* SK1 strain using forward primer 258 and reverse primer 259b. The reverse primer introduced the sequence for the C-terminal STREP affinity tag. The amplified product was digested with *Bam*HI and *Hin*dIII (New England Biolabs) and cloned into corresponding sites of pFB-GST-MLH1^18^ to create pFB-yMSH4-STREP. The yeast *MSH5* gene was amplified from the genomic DNA of the *S. cerevisiae* W303 strain using forward primer 265 and reverse primer 266. The *MSH5* gene was then cloned into *Bam*HI and *Xho*I restriction sites of pFB-MBP-MLH3-HIS^18^ to create pFB-yMSH5-HIS.

### Purification of hMLH1-hMLH3

The bacmids and baculoviruses were prepared individually using pFB-FLAG-hMLH1co and pFB-HIS-MBP-hMLH3co vectors according to manufacturer’s instructions (Bac-to-Bac system, Life Technologies). *Spodoptera frugiperda* 9 (*Sf*9) cells were seeded at 500,000 cells per ml 16 h before infection. The cells were then co-infected with both baculoviruses and incubated for 52 h at 27 °C with constant agitation. The cells were then harvested (500 x g, 10 min) and washed once with PBS (137 mM NaCl, 2.7 mM KCl, 10 mM Na_2_HPO_4_, 1.8 mM KH_2_PO_4_). The pellets were snap-frozen in liquid nitrogen and stored at −80 °C. All subsequent steps were carried out on ice or at 4 °C. The pellets were resuspended in 3 volumes of lysis buffer [50 mM Tris-HCl pH 7.5, 1 mM dithiothreitol (DTT), 1 mM ethylenediaminetetraacetic acid (EDTA), 1 mM phenylmethylsulfonyl fluoride (PMSF), 1:400 (volume/volume) protease inhibitor cocktail (Sigma, P8340), 30 μg/ml leupeptin (Merck)] and incubated for 20 min with continuous stirring. Next, 1/2 volume of 50% glycerol was added, followed by 6.5% volume of 5 M NaCl (final concentration 305 mM). The suspension was further incubated for 30 min with continuous stirring. The cell suspension was centrifuged for 30 min at 48,000 x g to obtain soluble extract. The supernatant was transferred to tubes containing pre-equilibrated Amylose resin (New England Biolabs, 4 ml per 1l of *Sf*9 culture) and incubated for 1 h with continuous agitation. The resin was collected by spinning at 2,000 x g for 2 min and washed extensively batchwise and on a disposable column (10 ml, Thermo Fisher) with Amylose wash buffer [50 mM Tris-HCl pH 7.5, 1 mM β-mercaptoethanol (β-ME), 1 mM PMSF, 10% glycerol, 300 mM NaCl]. Protein was eluted with Amylose elution buffer [50 mM Tris-HCl pH 7.5, 0.5 β-ME, 1 mM PMSF, 10% glycerol, 300 mM NaCl, 10 mM maltose (Sigma)] and the total protein concentration was estimated by Bradford assay. To cleave off the maltose binding tag (MBP), 1/6 (weight/weight) of PreScission protease (PP)^47^, with respect to total protein concentration in the eluate, was added and incubated for 1 h. Next, the cleaved amylose eluate was diluted by adding 1/2 volume of FLAG dilution buffer (50 mM Tris-HCl pH 7.5, 1 mM PMSF, 10% glycerol, 300 mM NaCl) to lower the concentration of β-ME. The diluted eluate was then incubated batchwise for 1 h with pre-equilibrated anti-FLAG M2 affinity resin (Sigma, A2220, 0.8 ml). The resin was washed extensively with FLAG wash buffer (50 mM Tris-HCl pH 7.5, 0.5 mM β-ME, 1 mM PMSF, 10% glycerol, 150 mM NaCl). Protein was eluted with FLAG wash buffer containing 150 ng/μl 3x FLAG peptide (Sigma), aliquoted, frozen in liquid nitrogen and stored at −80 °C. The final construct contained a FLAG tag at the N-terminus of hMLH1. The yield from 1 l culture was ∼0.5 mg and the concentration ∼2 μM. All hMLH1-hMLH3 mutants were expressed and purified using the same procedure.

### Purification of hMSH4-hMSH5

The human hMSH4-hMSH5 complex was expressed from a dual pFB-hMSH4co-STREP-hMSH5co-HIS vector in *Sf*9 cells using the Bac-to-Bac system as described above. All purification steps were carried out on ice or at 4 °C. The cell pellets were resuspended in 3 volumes of nickel-nitriloacetic acid (NiNTA) lysis buffer [50 mM Tris-HCl pH 7.5, 2 mM β-ME, 1 mM EDTA, 1 mM PMSF, 1:400 (volume/volume) protease inhibitor cocktail (Sigma, P8340), 30 μg/ml leupeptin (Merck), 20 mM imidazole] and incubated for 20 min with continuous stirring. Next 1/2 volume of 50% glycerol was added, followed by 6.5% volume of 5 M NaCl (final concentration 305 mM), and the suspension was further incubated for 30 min with continuous stirring. To obtain soluble extract, the suspension was centrifuged at 48,000 x g for 30 min. The soluble extract was transferred to a tube containing pre-equilibrated NiNTA resin (Qiagen, 4 ml per 1 l *Sf*9 cells) and incubated for 1 h with continuous mixing. The NiNTA resin was collected by centrifugation at 2,000 x g for 2 min. The resin was washed extensively batchwise and on a disposable column with NiNTA wash buffer (50 mM Tris-HCl pH 7.5, 2 mM β-ME, 300 mM NaCl, 1 mM PMSF, 10% glycerol, 20 mM imidazole). Protein was eluted with NiNTA wash buffer containing 250 mM imidazole. The eluted sample was incubated with pre-equilibrated Strep-Tactin Superflow resin (Qiagen, 0.7 ml) for 90 min with continuous mixing. The resin was transferred to a disposable column and washed extensively with Strep wash buffer (50 mM Tris-HCl pH 7.5, 2 mM β-ME, 300 mM NaCl, 1 mM PMSF, 10% glycerol). Protein was eluted with Strep wash buffer containing 2.5 mM d-Desthiobiotin (Sigma) and stored at −80°C after snap freezing in liquid nitrogen. The final construct contained a STREP tag at the C-terminus of hMSH4 and a HIS-tag at the C-terminus of hMSH5. The variants of the hMSH4-hMSH5 complex were purified using the same procedure. We note that the double mutant hMSH4G685A-hMSH5G597A heterodimer was not stable and could not be purified.

### Purification of hEXO1(D173A)

The pFB-EXO1(DA)-FLAG vector was used to prepare recombinant baculovirus and the protein was expressed in *Sf*9 cells as described above. Frozen cell pellet was thawed and resuspended in 3 pellet volumes of lysis buffer [50 mM Tris-HCl pH 7.5, 0.5 mM β-ME, 1 mM EDTA, 1:400 (volume/volume) protease inhibitor cocktail (Sigma, P8340), 0.5 mM PMSF, 20 μg/ml leupeptin]. The cell suspension was incubated with gentle stirring for 10 min. 1/2 volume of 50% glycerol and 6.5% volume of 5 M NaCl (final concentration 305 mM) were added. The suspension was incubated for 30 min with stirring. The extract was then centrifuged at 48,000 x g for 30 min. The soluble extract was added to pre-equilibrated M2 anti FLAG affinity resin (Sigma, A2220, 2 ml resin for purification from 1 l *Sf*9 cell culture) and incubated batchwise for 45 min. The suspension was then centrifuged (2,000 x g, 5 min), the supernatant (FLAG flowthrough) removed, and the resin was transferred to a disposable chromatography column. The resin was washed with 50 resin volumes of TBS buffer (20 mM Tris-HCl pH 7.5, 150 mM NaCl, 0.5 mM β-ME, 0.5 mM PMSF, 10% glycerol) supplemented with 0.1% NP40. This was followed by washing with 10 resin volumes of TBS buffer without NP40. EXO1-FLAG was eluted with TBS buffer supplemented with 150 ng/μl 3x FLAG peptide (Sigma, F4799). Fractions containing detectable protein (as estimated by the Bradford method) were pooled, applied on a disposable column with 1 ml pre-equilibrated Biorex70 resin (Bio-Rad), and flow-through was collected. The sample was then diluted by adding 1 volume of dilution buffer (50 mM Tris-HCl pH 7.5, 5 mM β-ME, 0.5 mM PMSF, 10% glycerol). Diluted FLAG-EXO1 was applied on 1 ml HiTrap SP HP column (GE Healthcare) pre-equilibrated with S buffer A (50 mM Tris-HCl pH 7.5, 75 mM NaCl, 5 mM *β*-ME, 10% glycerol) at 0.8 ml/min. The column was washed with 20 ml S buffer A, and eluted with 8 ml linear salt gradient in S buffer A (75 mM to 1 M NaCl). Peak fractions were pooled, aliquoted, frozen in liquid nitrogen and stored at −80 °C. The procedure yielded around ∼0.15 mg of protein from 1 l of *Sf*9 culture, with an approximate concentration of ∼1 μM.

### Purification of hMSH2-hMSH6 and hMSH2-hMSH3 heterodimers

To prepare the hMSH2-hMSH6 heterodimer, the *Sf*9 cells were co-infected with recombinant baculoviruses prepared from pFB-hMSH2-FLAG and pFB-hMSH6-HIS^46^ vectors. The purification was carried out at 4 °C or on ice. The cell pellets were resuspended in 3 volumes of lysis buffer [50 mM Tris-HCl pH 7.5, 1:400 [volume/volume] protease inhibitor cocktail (Sigma, P8340), 1 mM PMSF, 60 μg/ml leupeptin and 0.5 mM β-ME]. The sample was incubated while stirring for 20 min. 1/2 volume of 50% glycerol was added, followed by 6.5% volume 5 M NaCl (final concentration 305 mM). The cell suspension was incubated for 30 min with stirring. To obtain soluble extract, the suspension was centrifuged (30 min, 48,000 x g). The supernatant was mixed with pre-equilibrated 2 ml NiNTA resin (purification from 800 ml *Sf*9 cells) and incubated batchwise for 1 h. The resin was then washed batchwise and on column with wash buffer [30 mM Tris-HCl pH 7.5, 1:1,000 (volume/volume) protease inhibitor cocktail (Sigma, P8340), 15 μg/ml leupeptin, 0.5 mM β-ME, 0.5 mM PMSF, 20 mM imidazole, 300 mM NaCl, 10% glycerol]. Bound protein was eluted with elution buffer [30 mM Tris-HCl pH 7.5, 1:1,000 (volume/volume) protease inhibitor cocktail (Sigma, P8340), 15 μg/ml leupeptin, 0.5 mM β-ME, 0.5 mM PMSF, 300 mM imidazole, 150 mM NaCl, 10% glycerol]. The pooled fractions were diluted with 7 volumes of dilution buffer (30 mM Tris-HCl pH 7.5, 15 μg/ml leupeptin, 0.5 mM β-ME, 0.5 mM PMSF, 150 mM NaCl, 10% glycerol), and mixed with 0.7 ml pre-equilibrated anti-FLAG M2 affinity gel (Sigma). The suspension was incubated batchwise for 60 min. The sample was centrifuged (5 min, 1,000 g) and resin was transferred to a disposable chromatography column. The resin was then washed extensively with dilution buffer. The heterodimer was eluted with dilution buffer supplemented with 200 μg/ml 3x FLAG peptide (Sigma). Eluates containing protein were pooled, aliquoted, frozen in liquid nitrogen and stored at −80 °C. The hMSH2-hMSH3 heterodimer was prepared using the same procedure, using pFB-hMSH3-HIS^48^.

### Purification of yMsh4-yMsh5

Baculoviruses expressing yMsh4 and yMsh5 were prepared individually using the Bac-to-Bac system and pFB-yMSH4-STREP and pFB-yMSH5-HIS vectors. *Sf*9 cells were co-infected with optimized ratios of both viruses to express both proteins together as a heterodimer. The cells were harvested 52 h after infection, washed with PBS, and the pellets were frozen in liquid nitrogen and stored at −80 °C until use. The subsequent steps were carried out on ice or at 4 °C. The cell pellet was resuspended in lysis buffer [50 mM Tris-HCl pH 7.5, 2 mM β-ME, 1 mM EDTA, 1:400 (volume/volume) protease inhibitor cocktail (Sigma, P8340), 1 mM PMSF, 30 μg/ml leupeptin, 20 mM imidazole)] for 20 min. Then, 50% glycerol was added to a final concentration of 16%, followed by 5 M NaCl to a final concentration of 305 mM. The suspension was incubated for further 30 min with gentle agitation. The total cell extract was centrifuged at 48,000 x g for 30 min to obtain soluble extract. The extract was then bound to NiNTA resin (Qiagen) for 60 min batchwise followed by extensive washing with NiNTA wash buffer (50 mM Tris-HCl pH 7.5, 2 mM β-ME, 300 mM NaCl, 10 % glycerol, 1 mM PMSF, 10 μg/ml leupeptin, 20 mM imidazole) both batchwise and on a column. The heterodimer was eluted by NiNTA elution buffer (NiNTA wash buffer containing 250 mM imidazole). The eluate was further incubated with pre-equilibrated Strep-Tactin Superflow resin (Qiagen) for 60 min batchwise. The protein-bound resin was then washed in two sequential steps; first with STREP wash buffer I (50 mM Tris pH 7.5, 2 mM β-ME, 10 % glycerol, 1 mM PMSF and 300 mM NaCl) and then with STREP wash buffer II (50 mM Tris pH 7.5, 2 mM β-ME, 10 % glycerol, 1 mM PMSF and 50 mM NaCl). The heterodimer was eluted with STREP wash buffer II containing 2.5 mM d-Desthiobiotin (Sigma). The eluate was then applied on a pre-equilibrated HiTrap Q HP column (GE Healthcare). The column was washed with STREP wash buffer II and protein was eluted with a linear gradient of NaCl (50 to 600 mM) in STREP wash buffer II. Collected fractions were analyzed on SDS-PAGE, peak samples were pooled, aliquoted and stored at −80 °C. The final construct contained a STREP tag at the C-terminus of yMsh4 and a HIS-tag at the C-terminus of yMsh5. The procedure yielded ∼ 0.15 mg of protein from 4 l of *Sf*9 culture, with an approximate concentration of ∼ 1 μM.

### Purification of yMlh1-yMlh3

The yMlh1-yMlh3 heterodimer was expressed using pFB-HIS-yMLH1 and pFB-MBP-yMLH3 and the Bac-to-Bac system and purified using affinity chromatography^18^. Briefly, the cells were resuspended in lysis buffer containing 50 mM Tris-HCl pH 7.5, 1 mM DTT, 1 mM EDTA, 1:400 (volume/volume), protease inhibitor cocktail (Sigma, P8340), 1 mM PMSF, 30 μg/ml leupeptin and incubated for 20 min. Subsequently glycerol [final concentration 16% (volume/volume)] and NaCl (final concentration 305 mM) were added. Upon further incubation for 30 min and centrifugation (48,000 x g, 30 min), the cleared extract was then subjected to affinity chromatography with Amylose resin (New England Biolabs), the MBP tag was cleaved with PreScission protease and the heterodimer was further purified on Ni-NTA agarose (Qiagen)^18^. The final eluate was dialyzed into 50 mM Tris-HCl pH 7.5, 5 mM β-ME, 10% glycerol, 0.5 mM PMSF and 300 mM NaCl. Aliquots were flash frozen and stored at −80 °C until use. The purification yielded ∼ 1 mg protein from 2.4 l culture and the concentration was 5.9 μM.

### Purification of RFC, PCNA and the Ku heterodimer

Human PCNA was expressed in *E. coli* cells (1 l) from pET23C-his-hPCNA vector (a kind gift from Ulrich Huebscher, University of Zurich). Transformed cells were grown to OD 0.5, and induced with 0.5 mM isopropyl β-D-1-thiogalactopyranoside (IPTG) for 3.5 h at 37 °C. Cells were lysed by sonication in lysis buffer (20 mM Tris-HCl pH 7.5, 250 mM NaCl, 2 mM β-ME, 5 mM imidazole, 1 mM PMSF, 1:250 Sigma protease inhibitor cocktail P8340). The lysate was cleared by centrifugation (48,000 x g, 30 min) and bound to 2 ml NiNTA resin (Qiagen) for 1 h batchwise. Resin was washed with wash buffer (20 mM Tris-HCl pH 7.5, 250 mM NaCl, 2 mM β-ME, 30 mM imidazole, 1 mM PMSF), and PCNA was eluted with elution buffer (wash buffer supplemented with 400 mM imidazole). The sample was diluted to conductivity corresponding to 100 mM NaCl, and loaded on HiTrapQ column. The column was developed by a salt gradient (100 mM to 1 M NaCl) in 20 mM Tris-HCl pH 7.5, 2 mM β-ME and 10% glycerol. The fractions containing PCNA were pooled, aliquoted and stored at −80°C.

Yeast RFC was expressed in *E. coli* cells (4 l) transformed with pEAO271 (a kind gift from E. Alani, Cornell university). Cells were grown to OD 0.5, and induced with 0.5 mM isopropyl β-D-1-thiogalactopyranoside (IPTG) for 3 h at 37 °C. Cells were resuspended in lysis buffer (60 mM HEPES-NaOH pH 7.5, 250 mM NaCl, 2 mM β-ME, 0.5 mM EDTA, 1:250 Sigma protease inhibitor cocktail P8340, 1 mM PMSF, 10% glycerol) and disrupted by sonication. The cleared extract was loaded on 5 ml SP sepharose column, washed with buffer SP A (30 mM HEPES-NaOH pH 7.5, 300 mM NaCl, 2 mM β-ME, 0.5 mM EDTA, 1 mM PMSF, 10% glycerol) and eluted with a salt gradient (300 mM to 600 mM NaCl). Eluted fractions were analyzed by polyacrylamide gel electrophoresis, pooled and diluted to conductivity corresponding to 110 mM NaCl. The diluted sample was applied on HiTrapQ column, and eluted in 110 to 600 mM NaCl gradient in 30 mM HEPES-NaOH pH 7.5, 2 mM β-ME, 1 mM PMSF and 10% glycerol. The eluate was aliquoted and stored at −80 °C. The preparation of the yeast Ku heterodimer was described previously^49^.

### Nuclease assays

The reactions (15 μl) were carried out in 25 mM Tris-acetate pH 7.5, 1 mM DTT, 0.1 mg/ml bovine serum albumin (BSA, New England Biolabs), and as indicated manganese or magnesium acetate (5 mM), ATP (concentrations as indicated, GE Healthcare, 27-1006-01) and plasmid-based DNA substrate [100 ng per reaction, either 2.7 kbp-long pUC19 (Fig. 1), 5.6 kbp-long pFB-Rfa2 (Fig. 2-5), pAG25 (Addgene) or cruciform pIRbke8 mut^36^], where unlabeled DNA was used, or 1 nM, in molecules, where ^32^P-labeled substrate or detection method was used). Where indicated, ADP (Alfa Aesar, J60672), AMP-PNP (Sigma, A2647) or ATP-γ-S (Cayman, 14957) were used instead of ATP. Where indicated, the reactions were supplemented with PIP box peptide derived from p21 (GRKRRQTSMTDFYHSKRRLIFS) or control peptide with key residues mutated (underlined, GRKRR<u>A</u>TS<u>A</u>TDFYHSKRRLIFS). The reaction buffer was assembled on ice, and the recombinant proteins were then added on ice (hMLH1-hMLH3 protein was always added last). The reactions, unless indicated otherwise, were incubated for 60 min at 30 °C or 37 °C. The reactions were supplemented with protein storage or dilution buffer to compensate for components introduced with recombinant proteins in each particular experiment, this resulted in final NaCl concentrations ∼30 mM. The reactions were terminated with 5 μl STOP solution (150 mM EDTA, 2% SDS, 30% glycerol, 0.01% bromophenol blue), 1 μl proteinase K (Roche, 03115828001, 18 mg/ml) and further incubated for 60 min at 50 °C. The reaction products were then separated by electrophoresis in 1% agarose (Sigma, A9539) containing GelRed (Biotium) in TAE buffer. Using Bio-Rad SubCell GT system (gel length 26 cm), the separation was carried out for 90 min at 120 V. The gels were then imaged (InGenius3, GeneSys). The results were quantitated using ImageJ and expressed as % of nicked DNA versus the total DNA in each particular lane; any nicked DNA present in control (no protein) reactions was removed as a background.

### Electrophoretic mobility shift assays

The DNA binding reactions were carried out in 15 μl volume in binding buffer containing 25 mM HEPES pH 7.8, 5 mM magnesium chloride, 5% (volume/volume) glycerol, 1 mM DTT, 50 μg/ml BSA, 6.6 ng/μl dsDNA (competitor, 50 bp-long, 100-fold molar excess over labeled DNA), 0.5 nM DNA substrate (^32^P-labelled, in molecules) and respective concentrations of recombinant proteins (yeast or human MSH4-MSH5 complex and their variants, hMLH1-hMLH3 and variants, hEXO1). The oligonucleotide-based DNA substrates were ssDNA (labelled oligonucleotide PC1253), dsDNA (labelled PC1253 and PC1253C), Y-structure (labelled PC1254 and PC1253), HJ (labelled PC1253 and PC1254, PC1255 and PC1256) and D-Loop (labelled BB, and BT, INVa and INVb). MgCl_2_ was replaced by 3 mM EDTA where indicated. The reactions were assembled and incubated on ice for 15 min, followed by the addition of 5 μl EMSA loading dye (50% glycerol, 0.01% bromophenol blue). The products were separated on 6% native polyacrylamide gel (19:1 acrylamide-bisacrylamide, BioRad) on ice. The gels were dried on 17 CHR paper (Whatman), exposed to storage phosphor screens (GE Healthcare), and scanned by Phosphorimager (Typhoon FLA 9500, GE Healthcare). The quantitation was carried out by ImageQuant software (GE Healthcare) and graphs were plotted using Prism software (Prism 8, Graphpad). For the “super-shift” assays comprising yMlh1-yMlh3 and yMsh4-yMsh5, the reactions were carried out as mentioned above (with magnesium or EDTA, as indicated), except that the products were separated on 0.6% agarose gel in TAE buffer at 4 °C (1 h, 100 V). The gels were dried on DE81 paper (Whatman) and scanned as above. In the super-shift assays with hMLH1-hMLH3, hMSH4-hMSH5 and hEXO1, the reaction buffer additionally contained 75 mM NaCl and 10 μM ATP. The DNA binding assays with yKu70-80 were carried out similarly, without salt and ATP, and were incubated for 30 min at 30 °C.

### Protein interaction assays

To test for protein-protein interactions, recombinant “bait” protein was immobilized on beads coupled to a specific antibody and incubated with the “prey” protein. After removal of unbound protein by beads washing, proteins were either detected by silver staining or by western blot.

To test for the interaction between hMLH1-hMLH3 and hMSH4-hMSH5, 0.7 μg anti-MLH1 antibody (Abcam, ab92312) was captured on 15 μl Protein G magnetic beads (Dynabeads, Invitrogen) by incubating in 50 μl PBS-T (PBS with 0.02% Tween-20) for 60 min with gentle mixing at regular intervals. The beads were washed 3 times on magnetic racks with 150 μl PBS-T to remove unbound antibodies. The beads were then mixed with 165 nM recombinant hMLH1-hMLH3 and 220 nM hMSH4-hMSH5 in 50 μl binding buffer I (25 mM HEPES pH 7.8, 3 mM EDTA, 1 mM DTT, 50 μg/ml BSA, 54 mM NaCl) and incubated on ice for 45 min with gentle agitation at regular intervals. Beads were then washed 3 times with 150 μl wash buffer I (25 mM HEPES pH 7.8, 3 mM EDTA, 1 mM DTT, 0.02% Tween-20) and proteins were eluted by boiling the beads in SDS buffer (50 mM Tris-HCl pH 6.8, 1.6% sodium dodecyl sulphate, 100 mM DTT, 10% glycerol, 0.01% bromophenol blue) for 3 min at 95 °C. The eluate was separated on a 10% SDS-PAGE gel and proteins were detected by silver staining. To perform the experiment reciprocally, 5 μg anti-HIS antibody (Genscript, A00186) was captured on Protein G beads (Dynabeads, Invitrogen) as described above. The recombinant protein complexes, as above, were then added and incubated in 50 μl binding buffer II (25 mM HEPES pH 7.8, 3 mM EDTA, 1 mM DTT, 50 μg/ml BSA, 137 mM NaCl) for 45 min with gentle agitation at regular intervals. Beads were then washed 3 times with wash buffer II (25 mM HEPES pH 7.8, 3 mM EDTA, 1 mM DTT, 80 mM NaCl, 0.1% Triton X-100). The subsequent steps were carried out as described above. To test for species-specific interactions as shown in Extended Data Fig. 4f, the same procedure was followed except 100 nM of either hMSH4-hMSH5 or yMsh4-yMsh5 was incubated with 400 nM hMLH1-hMLH3. To test for the interaction between yeast yMlh1-yMlh3 and yMsh4-yMsh5, 10 μl Protein G beads were used to capture 1 μg anti-STREP antibody (Biorad, MCA2489). yMsh4-yMsh5 (120 nM) was incubated with the beads in 60 μl binding buffer III (25 mM Tris-HCl pH 7.5, 3 mM EDTA, 1 mM DTT, 20 mg/ml BSA, 68 mM NaCl) for 60 min with continuous mixing. Next, the beads were washed 3 times with 150 μl wash buffer III (25 mM Tris-HCl pH 7.5, 3 mM EDTA, 1 mM DTT, 120 mM NaCl, 0.05% Triton X-100). 300 nM yMlh1-yMlh3 was then added to the resuspended beads in 60 μl binding buffer III, and incubated for additional 60 min with continuous mixing. Beads were washed 3 times with 150 μl wash buffer III and boiled afterwards for 3 min at 95 °C in SDS buffer to elute the proteins. The protein complexes were detected by western blot with anti-HIS antibody (Genscript, A00186).

To test for the interaction between hMLH1-hMLH3 and hEXO1(D173A), 0.33 μg anti-MLH1 antibody (Abcam ab223844) was captured on 10 μl protein G magnetic beads (Dynabeads, Invitrogen) by incubating in 50 μl PBS-T (PBS with 0.1% Tween-20) for 2 h at 4 °C with gentle mixing at regular intervals. The beads were washed 4 times on magnetic racks with 150 μl PBS-T to remove unbound antibody. The beads were then mixed with 1 μg recombinant hMLH1-hMLH3 and 0.5 μg hEXO1(D173A) in 200 μl binding buffer I (25 mM Tris-HCl pH 7.5, 3 mM EDTA, 1 mM DTT, 20 μg/ml BSA, 300 mM NaCl) and incubated on ice for 2 h with gentle agitation at regular intervals. Beads were then washed 4 times with 300 μl wash buffer I (50 mM Tris-HCl pH 7.5, 3 mM EDTA, 1 mM DTT, 300 mM NaCl, 0.05% Triton X-100) and proteins were eluted by boiling the beads in SDS buffer (50 mM Tris-HCl pH 6.8, 1.6% sodium dodecyl sulphate, 100 mM DTT, 10% glycerol, 0.01% bromophenol blue) for 3 min at 95 °C. The eluate was separated on a 10% SDS-PAGE gel and proteins were detected by silver staining.

To test for the interaction between hMLH1-hMLH3 and hPCNA or hEXO1, 1 μg anti-MLH1 antibody (Abcam ab223844) was captured on 15 μl protein G magnetic beads (Dynabeads, Invitrogen) by incubating in 50 μl PBS-T (PBS with 0.1% Tween-20) for 1 h at room temperature with gentle mixing at regular intervals. The beads were washed 3 times on magnetic racks with 150 μl PBS-T to remove unbound antibody. The beads were then mixed with 1.5 μg each recombinant hMLH1-hMLH3 and hPCNA or hEXO1, in 60 μl binding buffer I (25 mM Tris-HCl pH 7.5, 3 mM EDTA, 1 mM DTT, 20 μg/ml BSA, 60 mM NaCl) and incubated on ice for 1 h with gentle agitation at regular intervals. Beads were then washed 4 times with 150 μl wash buffer I (50 mM Tris-HCl pH 7.5, 3 mM EDTA, 1 mM DTT, 120 mM NaCl, 0.05% Triton X-100) and proteins were eluted by boiling the beads in SDS buffer (50 mM Tris-HCl pH 6.8, 1.6% sodium dodecyl sulphate, 100 mM DTT, 10% glycerol, 0.01% bromophenol blue) for 3 min at 95 °C. Avidin (Sigma, A9275, 110 ng/μl) was added to the eluate as a stabilizer. The eluate was separated on a 10% SDS-PAGE gel and proteins were detected by silver staining.

### Yeast manipulations

All yeast strains are derivatives of the SK1 background and are listed in Table S1. Yeast strains were obtained by direct transformation or crossing to obtain the desired genotype. The following alleles have been described previously: *mlh1Δ*, *mlh3Δ* as well as spore-autonomous fluorescent marker for the live cell recombination assays^9, 50^.

YIplac211 plasmid derivatives carrying *MLH1* (pYIplac211-*MLH1*) or *MLH3* (pYIplac211-*MLH3*), as well as the respective promoter (∼ 500bp upstream of ATG) and terminator (∼ 200bp downstream of STOP) regions were used to complement *mlh1Δ* or *mlh3Δ* mutant strains, respectively. pYIplac211-*MLH1* and pYIplac211-*MLH3* were linearized and integrated in the promoter region of the respective genomic loci. pYIplac211-*MLH1^Q572A-L575A-F578A^* (encoding Mlh1^P^) and pYIplac211-*MLH3^Q293A-V296A-F300A^* (encoding Mlh3^P^) were generated by restriction digest-mediated insertion of a synthetic fragment carrying the respective mutations into pYIplac211-*MLH1* or pYIplac211*-MLH3*.

Rfc1 was C-terminally tagged with TAP tag. The Mlh1-HA allele was described previously^51^. Transformants were confirmed using PCR discriminating between correct and incorrect integrations and sequencing. All experiments were performed at 30 °C. Two different approaches were used for meiosis induction. In the first one, cells were grown in SPS presporulation medium and transferred in sporulation medium as described^52^. For highly synchronous copper-inducible meiosis, the procedure as described^53^. Briefly, cells were grown in YPD to exponential phase. Exponentially growing yeast were inoculated at OD_600_ = 0.05 into reduced glucose YPD (1% yeast extract, 2% peptone, 1% glucose) and grown to an OD_600_ = 11-12 for 16-18 h. Cells were washed, resuspended in sporulation medium (1.0% [w/v] potassium acetate, 0.02% [w/v] raffinose, 0.001% polypropylene glycol) at OD_600_ = 2.5. After 2 h, copper(II) sulfate (50 µM) was added to induce *IME1* expression from the *CUP1* promoter.

### Analysis of recombination using spore-autonomous fluorescence

The spore-autonomous fluorescence analysis of recombination was performed as described^50^, with some minor modifications. Diploid yeast cell colonies were streaked on YP_2%glycerol_ plates, grown for 48 h, and single colonies were expanded twice in YPD plates at 30 °C for 24 h. Cells were then transferred to sporulation medium plates (SPM, 2% KAc) and incubated at 30 °C for 48 h. Spores were resuspended in SPM, briefly sonicated and transferred onto Poly-L-Lysine coated microscopy slides. Images were captured in four channels using a Wide-field DeltaVision multiplexed microscope with a 60x 1.4NA DIC Oil PlanApoN objective and a peco.edge 5.5 camera under the control of Softworx (Applied Precision). Images were processed in Fiji and the pattern of spore fluorescence in tetrads was manually scored. Only tetrads with each fluorescent marker occurring in two spores were included in the final assay. Recombination frequency, expressed as map distance in centimorgans was calculated using the Stahl lab online tools (https://elizabethhousworth.com/StahlLabOnlineTools/) ^54^. Three biological replicates using independent clones were analyzed. ≥900 tetrads were scored for each genotype.

### Analysis of spore viability

Spore viability was determined by microdissection of ≥ 156 spores from at least two biological replicates after induction of meiosis on SPM plates at 30 °C for 24 h.

### Co-immunoprecipitation and Western blot analysis

1.2×10^9^ cells were harvested, washed once with PBS, and lyzed in 3 ml lysis buffer [20 mM HEPES-KOH pH 7.5, 150 mM NaCl, 0.5% Triton X-100, 10% glycerol, 1 mM MgCl_2_, 2 mM EDTA; 1 mM PMSF; 1 x Complete Mini EDTA-Free (Roche); 1X PhosSTOP (Roche); 125 U/ml benzonase] with glass beads three times for 30 s in a Fastprep instrument (MP Biomedicals, Santa Ana, CA). The lysate was incubated 1 h at 4 °C. 100 μl of PanMouse IgG magnetic beads (Thermo Scientific) were washed with 100 μl lysis buffer, preincubated in 100 μg/ml BSA in lysis buffer for 2 h at 4 °C and then washed twice with 100 μl lysis buffer. The lysate was cleared by centrifugation at 13,000 x g for 5 min and incubated overnight at 4 °C with washed PanMouse IgG magnetic beads. The magnetic beads were washed four times with 1 ml wash buffer [20 mM HEPES-KOH pH7.5, 150 mM NaCl, 0.5% Triton X-100, 5% Glycerol, 1 mM MgCl_2_, 2 mM EDTA, 1 mM PMSF, 1 x Complete Mini EDTA-Free (Roche)]. The beads were resuspended in 30 μl TEV-C buffer (20 mM Tris-HCl pH 8, 0.5 mM EDTA, 150 mM NaCl, 0.1% NP-40, 5% glycerol, 1 mM MgCl2, 1 mM DTT) with 3 μl TEV protease (1 mg/ml) and incubated for 2 h at 23°C under agitation. The eluate was transferred to a new tube. Beads eluate was heated at 95 °C for 10 min and loaded on polyacrylamide gel [4-12% Bis-Tris gel (Invitrogen)] and run in MOPS SDS Running Buffer (Life Technologies). Proteins were then transferred to PVDF membrane using Trans-Blot® Turbo™ Transfer System (Biorad) at 1 A constant, up to 25 V for 45 min. Proteins were detected using c-Myc mouse monoclonal antibody (9E10, Santa Cruz, 1:500), HA.11 mouse monoclonal antibody (16B12, Biolegend, 1/750) or TAP rabbit monoclonal antibody (Invitrogen, 1:4,000). The TAP antibody still detects the CBP (Calmodulin Binding Protein) moiety after TEV cleavage of the TAP tag. Signal was detected using the SuperSignal West Pico or Femto Chemiluminescent Substrate (ThermoFisher). Images were acquired with a Chemidoc system (Biorad).

### Chromatin immunoprecipitation, real-time quantitative PCR and ChIPseq

For each meiotic time point, 2×10^8^ cells were processed as described^55^, with the following modifications: lysis was performed in lysis buffer plus 1 mM PMSF, 50 μg/ml aprotinin and 1x Complete Mini EDTA-Free (Roche), using 0.5 mm zirconium/silica beads (Biospec Products, Bartlesville, OK). The lysate was directly applied on 50 μl PanMouse IgG magnetic beads. Before use, magnetic beads were blocked with 5 μg/μl BSA for 4 h at 4 °C. Quantitative PCR was performed from the immunoprecipitated DNA or the whole cell extract using a QuantStudio 5 Real-Time PCR System and SYBR Green PCR master mix (Applied Biosystems, Thermo Scientific) as described^55^. Results were expressed as % of DNA in the total input present in the immunoprecipitated sample and normalized by the negative control site in the middle of *NFT1*, a 3.5 kb long gene. For the meiotic time-course in Figure 5f, the data were further normalized by the value at the 2 h time-point (time of meiosis induction by copper addition). Primers for *GAT1*, *BUD23*, *HIS4LEU2*, Axis and *NFT1* have been described.

---

**Extended Data Figure 1.**
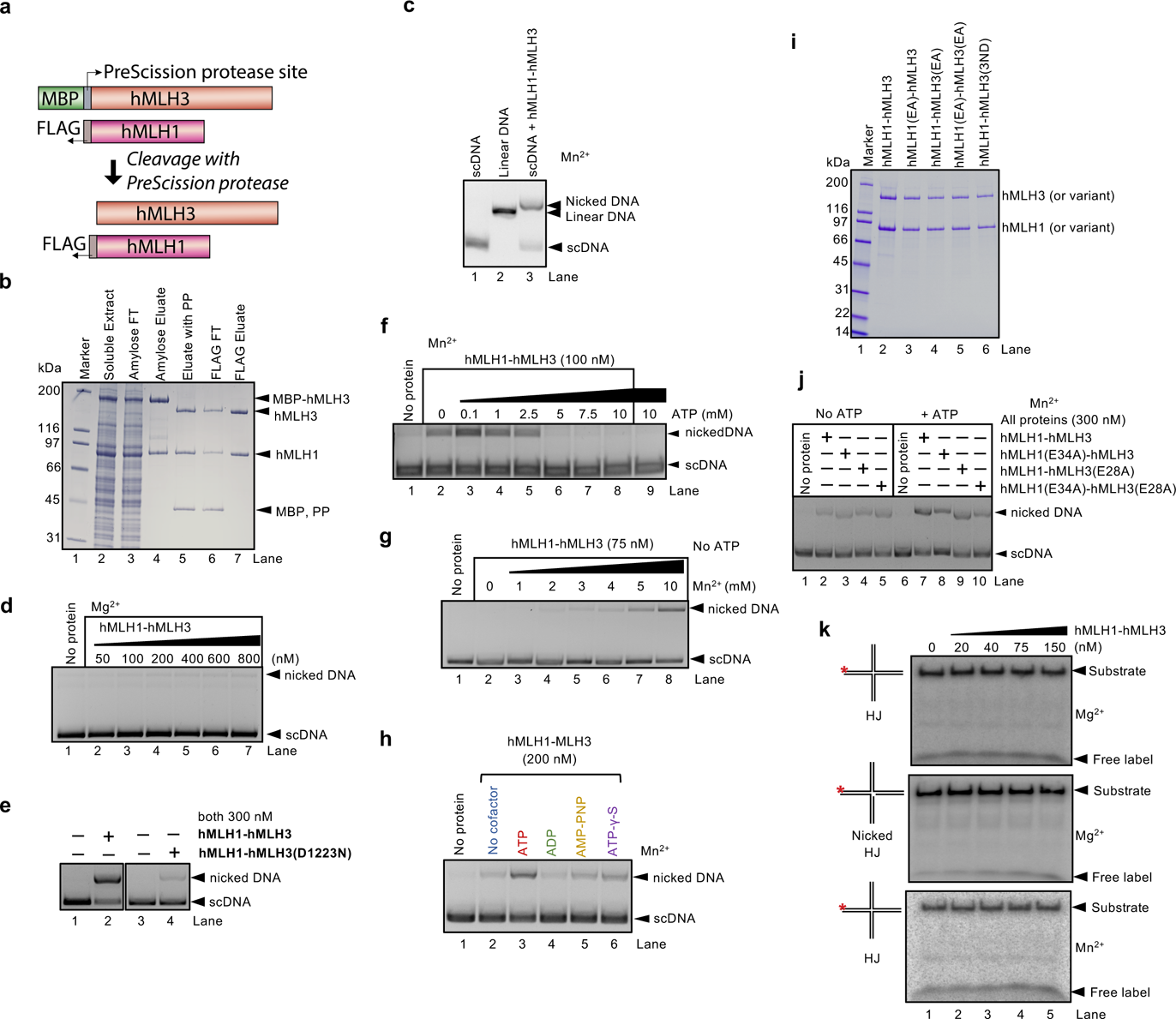
hMLH1-hMLH3 nicks scDNA. **a**, A scheme of hMLH1 and hMLH3 constructs. The maltose-binding protein (MBP) on hMLH3 was cleaved during protein purification. **b**, A representative purification of the hMLH1-hMLH3 complex. Amylose FT, flowthrough from Amylose resin; FLAG-FT, flowthrough from anti-FLAG resin; PP, PreScission protease; MBP, maltose-binding protein. The 4-15% gradient polyacrylamide gel was stained with Coomassie Blue. **c**, Nuclease assay with hMLH1-hMLH3 (300 nM) and pUC19 (2.6 kbp) scDNA substrates. Linear DNA was used a marker. The assay was carried out at 37 °C and contained 5 mM manganese acetate and ATP (0.5 mM). The hMLH1-hMLH3 nuclease introduces nicks in dsDNA but does not linearize dsDNA. **d**, Nuclease assay with hMLH1-hMLH3 and 5 mM magnesium acetate. The reaction buffer contained ATP (0.5 mM). The assay was carried out at 37 °C. The heterodimer exhibits barely detectable nuclease activity in magnesium. **e**, Representative nuclease assays with pUC19 dsDNA, with ATP (0.5 mM), and either wild type or hMLH1-hMLH3(D1223N), with a single amino acid substitution in the metal binding motif of hMLH3. The mutant retained ∼10% nuclease activity, and therefore a variant with three substitutions in the nuclease motif was used in this work (see Fig. 1d and further below). **f**, Nuclease assay with hMLH1-hMLH3 and manganese acetate in the presence of various concentrations of ATP, carried out at 37 °C. Low concentrations of ATP stimulated DNA cleavage, while elevated ATP concentrations were inhibitory. The inhibitory effect is likely due to a decrease in free manganese concentration (see panel g). **g**, Nuclease assay with hMLH1-hMLH3 and various concentrations of manganese acetate. The assay was carried out at 37 °C. **h**, Nuclease assay with hMLH1-hMLH3 and various cofactors (ADP, ATP and non-hydrolysable ATP analogs ATP-γ-S and AMP-PNP, all 0.5 mM). The assay was carried out at 37 °C with 5 mM manganese acetate. See Fig. 1f for quantitation of this and similar experiments. **i**, Purified hMLH1-hMLH3 variants used in this study. hMLH1(EA), hMLH1(E34A); hMLH3(EA), hMLH3(E28A); hMLH3(3ND), hMLH3 (D1223N, Q1224K, E1229K). **j**, Nuclease assay with wild type hMLH1-hMLH3 and indicated variants deficient in ATP hydrolysis, without or with ATP (0.5 mM). The assay was carried out at 37 °C, with 5 mM manganese acetate. See Fig. 1g for quantitation of this and similar experiments. **k**, Nuclease assays with wild type hMLH1-hMLH3 on oligonucleotide-based DNA substrates (Holliday junction, HJ and nicked Holliday junction, nicked HJ). The asterisk indicates the position of the radioactive label. The assay was carried out at 37 °C, with 5 mM manganese acetate or magnesium acetate, as indicated, with ATP (1 mM). The products were analyzed by 15% denaturing polyacrylamide gel electrophoresis.

**Extended Data Figure 2.**
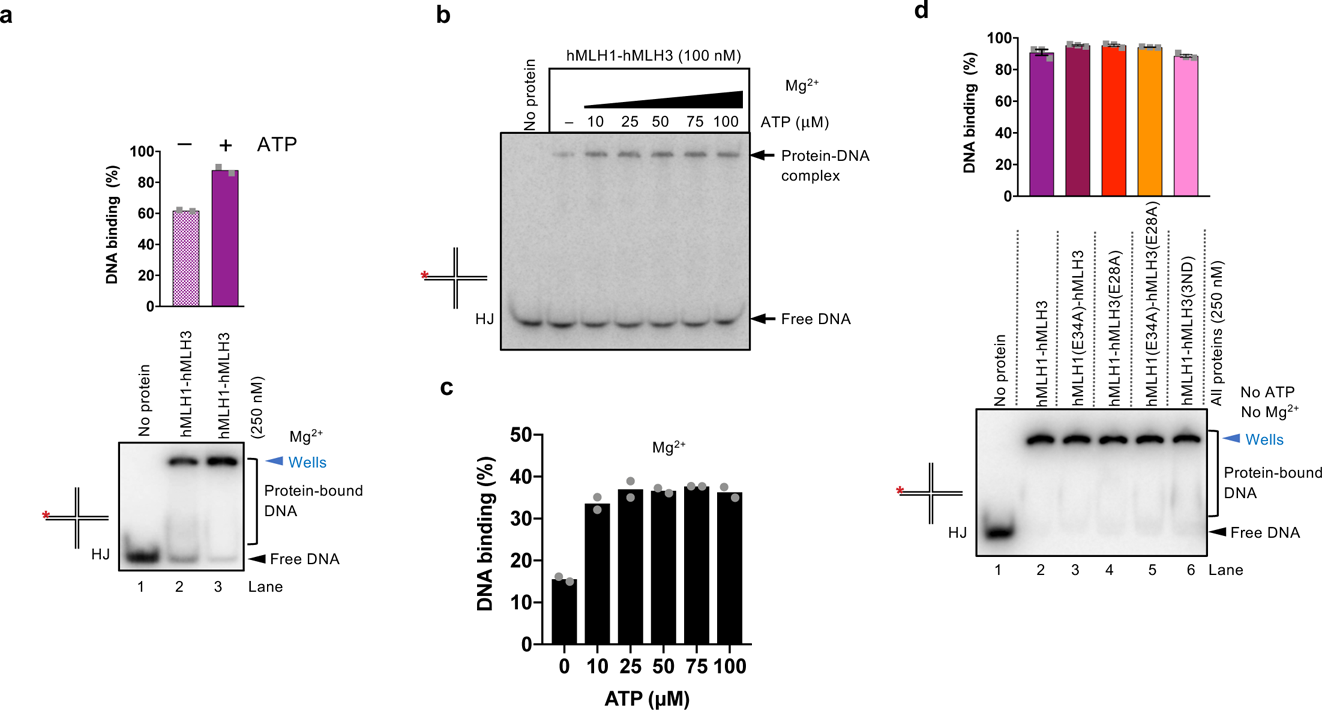
ATP promotes DNA binding by hMLH1-hMLH3. **a**, Electrophoretic mobility shift assay with hMLH1-hMLH3, without or with ATP (1 mM), with 2 mM magnesium acetate, using oligonucleotide-based HJ as the DNA substrate. Asterisk (*) indicates the position of the radioactive label. A representative experiment is shown at the bottom, a quantitation (average with individual values from two independent experiments) at the top. **b**, Electrophoretic mobility shift assay with hMLH1-hMLH3, oligonucleotide-based HJ as the DNA substrate, and various ATP concentrations, with 2 mM magnesium acetate, as indicated. The panel shows a representative experiment. **c**, Quantitation of experiments such as shown in panel b. The data points show averages and individual data points from two independent experiments. **d**, Electrophoretic mobility shift assay with indicated hMLH1-hMLH3 variants, oligonucleotide-based HJ as the substrate, in the absence of ATP and no magnesium (with 3 mM EDTA). Asterisk (*) indicates the position of the radioactive label. A representative experiment is shown at the bottom, a quantitation (averages shown, n=3; error bars, SEM) at the top.

**Extended Data Figure 3.**
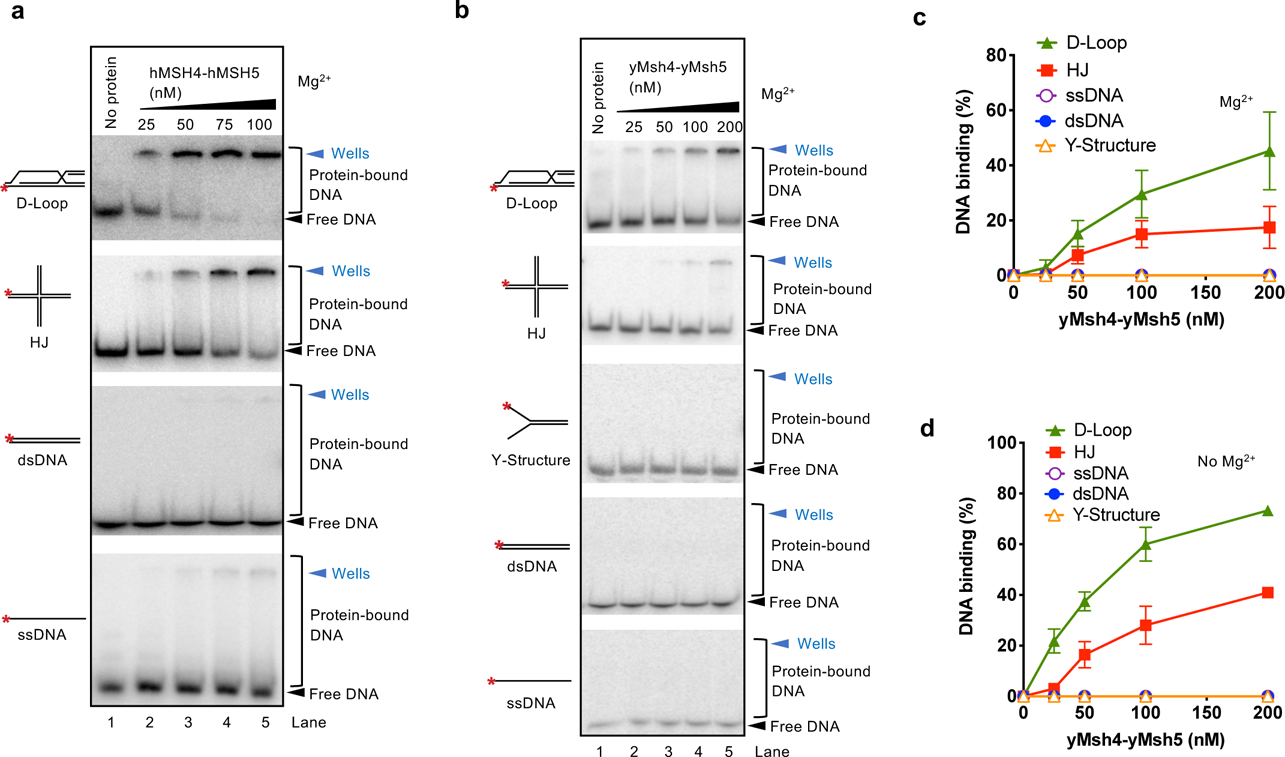
Human and yeast MutSγ complexes preferentially bind joint molecule DNA intermediates. **a**, Electrophoretic mobility shift assays with hMSH4-hMSH5 and indicated DNA substrates. Asterisk (*) indicates the position of the radioactive label. The assays were carried out in a buffer containing 2 mM magnesium acetate without ATP. For quantitation of this and similar experiments, see Fig. 2b. **b**, Electrophoretic mobility shift assays in 6% polyacrylamide gels with yMsh4-yMsh5 and indicated DNA substrates. Asterisk (*) indicates the position of the radioactive label. The assays were carried out in a buffer containing 2 mM magnesium acetate without ATP. **c**, Quantitation of experiments such as shown in panel b. Averages shown; error bars, SEM; n=3. **d**, Quantification of electrophoretic mobility shift assay with yMsh4-yMsh5 and indicated DNA substrates, without magnesium (with 3 mM EDTA). Averages shown; error bars, SEM; n=3.

**Extended Data Figure 4.**
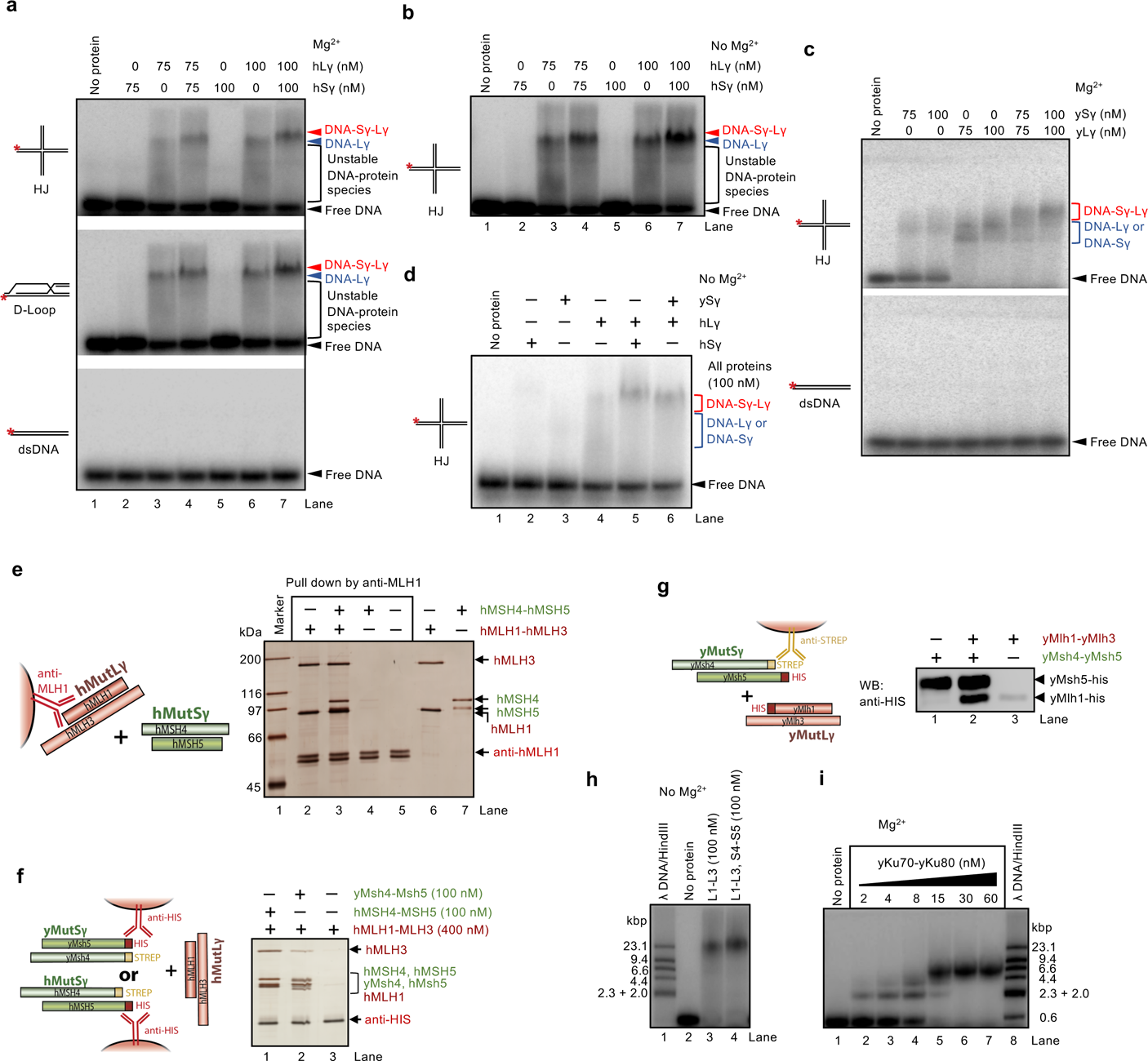
hMSH4-hMSH5 and hMLH1-hMLH3 interact and stabilize each other at DNA junctions. a, To investigate the interplay of hMutLγ and hMutSγ at DNA junctions, we performed electrophoretic mobility shift assays with either or both complexes under more stringent conditions (75 mM NaCl, 2 mM magnesium acetate), separated on 1% agarose gels. Here, hMSH4-hMSH5 lost the capacity to stably bind HJs/D-Loops, but could help stabilize the hMutSγ-hMutLγ complex. The binding of hMutLγ alone was not stable, as evidenced by a lack of a distinct protein-DNA band and the presence of smear in the lanes indicative of complexes that dissociated during electrophoresis. The addition of hMutSγ resulted in a moderate stabilization of the protein-DNA complex, and a minor super-shift in electrophoretic mobility. Shown are representative experiments. b, Electrophoretic mobility shift assays as in panel a, but without magnesium (with 3 mM EDTA). c, Electrophoretic mobility shift assays as in a, but with yeast MutLγ and MutSγ complexes. d, Assays as in a, with human hMLH1-hMLH3 and either human MSH4-MSH5 or yeast Msh4-Msh5. The supershift was observed only when the cognate human complexes were combined. e, Protein interaction assays with immobilized hMLH1-hMLH3 (bait) and hMSH4-hMSH5 (prey). The 10% polyacrylamide gel was stained with silver. f, Protein interaction assay with immobilized hMSH4-hMSH5 or yMsh4-yMsh5 that were used as baits, and hMLH1-hMLH3 (prey). The eluted proteins were analyzed by silver staining. Although residual interaction between yMsh4-yMsh5 and hMLH1-hMLH3 was still detected, it was much weaker than the interaction between the cognate hMSH4-hMSH5 and hMLH1-hMLH3 complexes. g, Protein interaction assay with immobilized yMsh4-yMsh5 (bait) and yMlh1-yMlh3 (prey). The eluted proteins were analyzed by western blotting. h, Electrophoretic mobility shift assays with hMLH1-hMLH3 (L1-L3) and hMSH4-hMSH5 (S4-S5), as indicated, and HJ DNA substrate. ^32^P-labeled λDNA/HindIII digest was used as a marker. The DNA-bound hMLH1-hMLH3 and hMSH4-hMSH5 species migrate high up on the agarose gel where the resolution capacity is limited. i, Electrophoretic mobility shift assay with yeast Ku70-Ku80 heterodimer and HJ DNA substrate. Ku bound the dsDNA ends of the four HJ arms, resulting in up to 4 heterodimers bound to the DNA substrate (lanes 5-7). Comparison with λ DNA/HindIII and panel h revealed that the Ku-DNA complex migrates much faster than DNA-bound hMLH1-hMLH3 and hMSH4-hMSH5. This suggests that multiple units of hMLH1-hMLH3 and hMSH4-hMSH5 bind DNA.

**Extended Data Figure 5.**
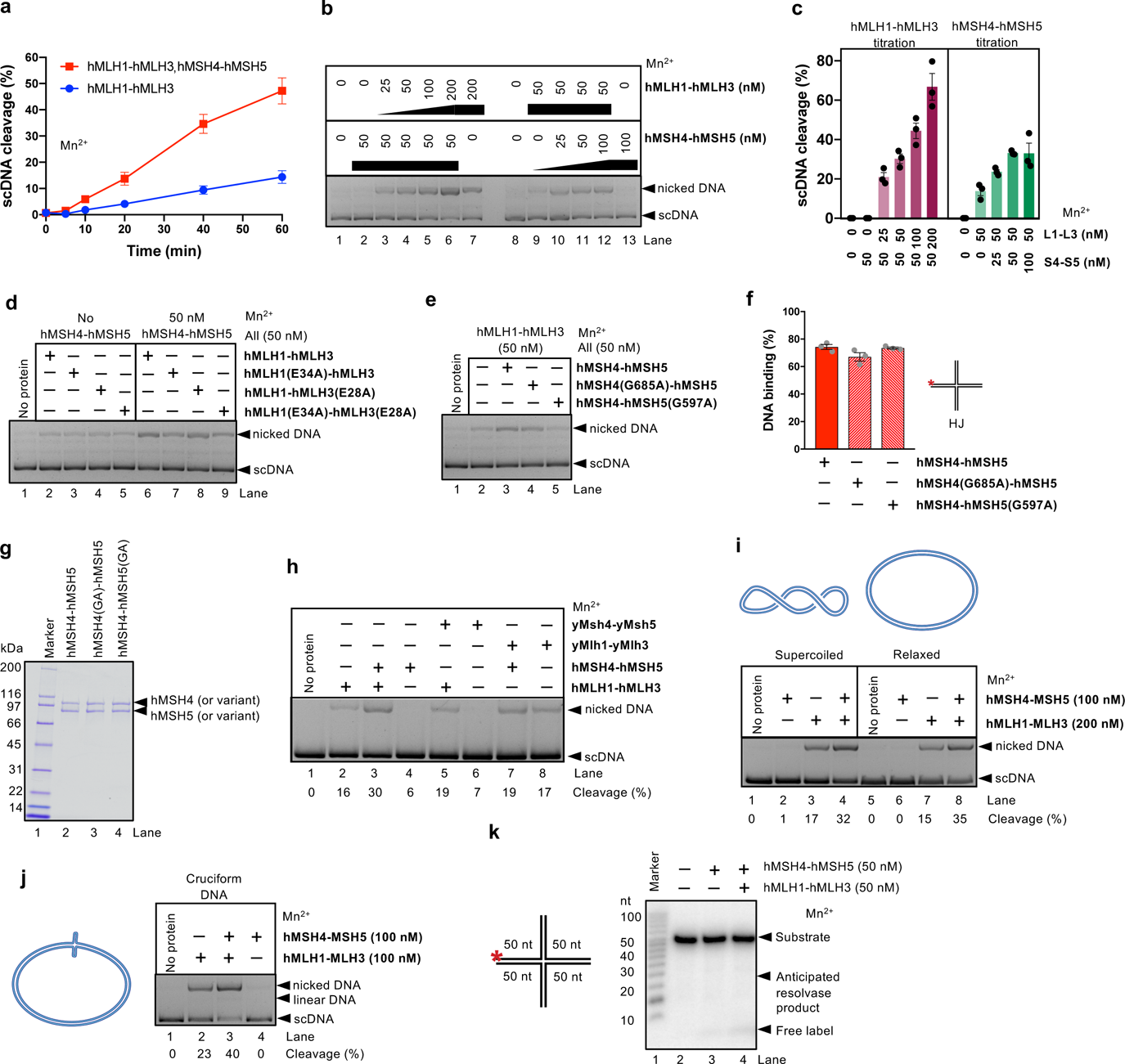
hMSH4-hMSH5 promotes DNA cleavage by hMLH1-MLH3, but does not exhibit resolvase activity. **a**, Quantitation of kinetic nuclease assays with hMLH1-hMLH3 (50 nM) without or with hMSH4-hMSH5 (50 nM). The assays were carried out at 30 °C in the presence of 5 mM manganese acetate and 2 mM ATP. Averages shown; error bars, SEM; n=3. **b**, Representative nuclease assays with various hMLH1-hMLH3 and hMSH4-hMSH5 concentrations, as indicated. The assays were carried out at 30°C in the presence of 5 mM manganese acetate and 0.5 mM ATP. **c**, Quantitation of experiments such as shown in panel b. Averages shown; error bars, SEM, n=3. The efficiency of nuclease cleavage was generally dependent on the concentrations used. When using 50 nM hMLH1-hMLH3, the maximal cleavage efficiency was achieved together with 50 nM hMSH4-hMSH5, no further increase when using 100 nM hMSH4-hMSH5 was observed. This suggests that both heterodimers may form a stoichiometric complex. *Vice versa*, when using 50 nM hMSH4-hMSH5, a further increase of DNA cleavage was observed when hMLH1-hMLH3 concentrations exceeded 50 nM, which is in agreement with the capacity of hMLH1-hMLH3 to cleave DNA on its own. **d**, Representative nuclease assays with hMSH4-hMSH5 and variants of hMLH1-hMLH3 deficient in ATP hydrolysis, as indicated. The assays were carried out at 30 °C in the presence of 5 mM manganese acetate and 0.5 mM ATP. For the quantification of these and similar data, see Fig. 2g. **e**, Representative nuclease assays with hMLH1-hMLH3 and variants of hMSH4-hMSH5 deficient in ATP hydrolysis, as indicated. The assays were carried out at 30 °C in the presence of 5 mM manganese acetate and 0.5 mM ATP. For the quantification of these and similar data, see Fig. 2h. **f**, Quantitation of electrophoretic mobility shift assays with hMSH4-hMSH5 and its ATPase motif mutant variants. Oligonucleotide-based HJ was used as the substrate. ATP was not included in the binding buffer. The mutations did not affect the capacity of hMSH4-hMSH5 to bind DNA. Averages shown; error bars, SEM; n=3. **g**, Recombinant hMSH4-hMSH5 and its variants used in this study. **h**, Nuclease reactions were carried out with yeast or human MutSγ and MutLγ complexes, as indicated (50 nM). While human MutSγ promoted DNA cleavage by MutLγ (compare lanes 2 and 3), yeast MutSγ did not promote DNA cleavage by human MutLγ (compare lanes 2 and 5), and reciprocally, human MutSγ did not promote DNA cleavage by yeast MutLγ (compare lanes 7 and 8). Continued on next page. Nuclease assay with supercoiled and relaxed DNA and recombinant proteins, as indicated. The MutSγ and MutLγ complexes cleave supercoiled and relaxed DNA with comparable efficiencies. The quantitation below the lanes represents an average from two independent experiments. j, Cleavage of pIRbke8mut cruciform DNA (inverted repeats folding back to form a Holliday junction structure) by MutSγ and MutLγ complexes. The quantitation below the lanes represents an average from two independent experiments. Simultaneous cleavage of both strands at the junction point would lead to linear DNA. No linear DNA was observed with MutSγ and MutLγ, indicating a lack of canonical resolvase activity. k, A representative nuclease assay with indicated proteins and oligonucleotide-based HJ DNA. No DNA cleavage was observed, indicating a lack of structure-specific DNA cleavage activity on the oligonucleotide-based substrate.

**Extended Data Figure 6.**
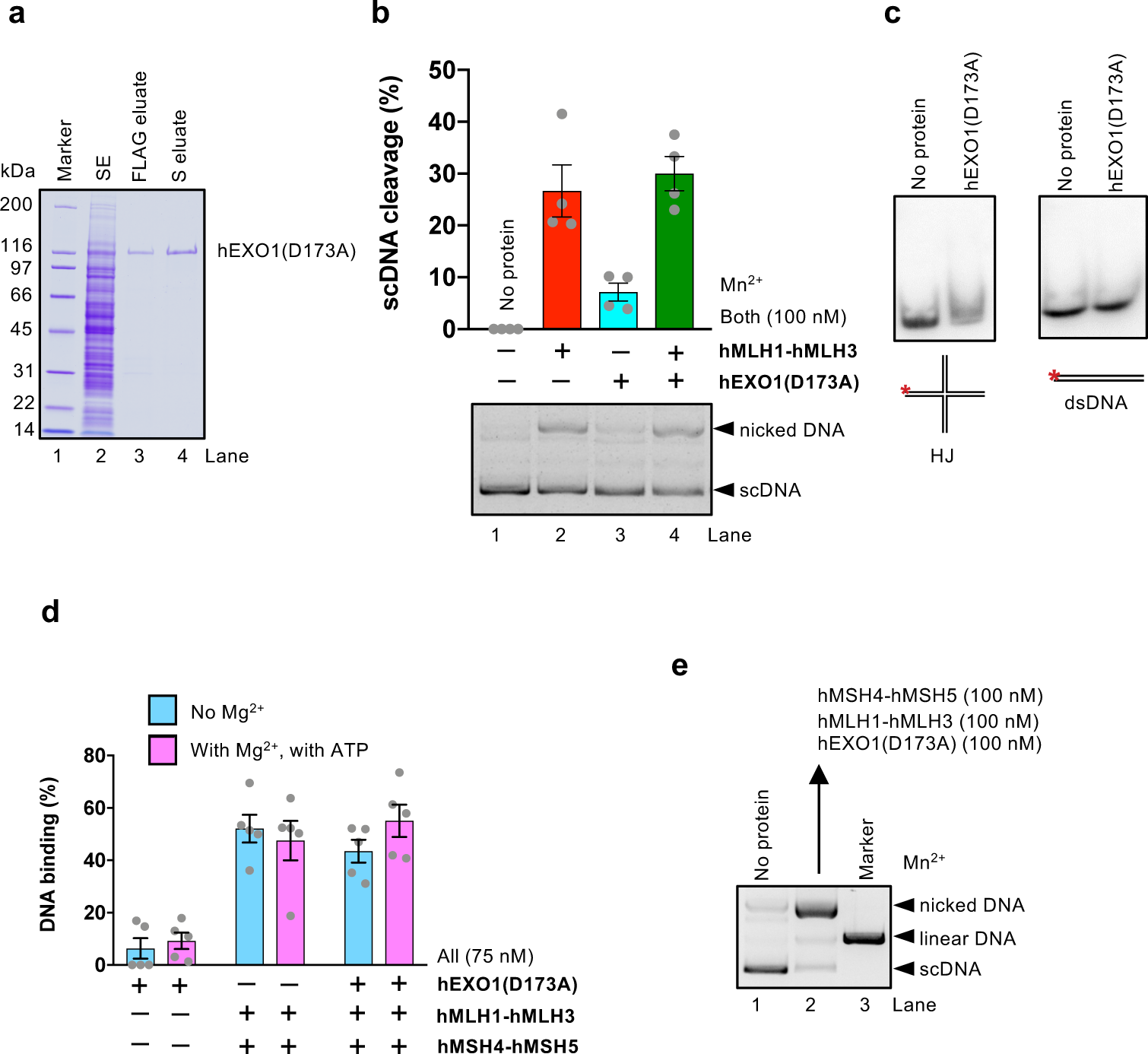
Stimulation of hMLH1-hMLH3 and hMSH4-hMSH5 by hEXO1(D173A). **a**, Purification of hEXO1(D173A). SE, soluble extract; FLAG eluate, eluate from anti-FLAG affinity resin; S eluate, eluate from HiTrap SP HP column. **b**, Nuclease assays with hMLH1-hMLH3 and/or hEXO1(D173A), as indicated. The assays were carried out at 30 °C in the presence of 5 mM manganese acetate and 0.5 mM ATP. A representative experiment is shown at the bottom, a quantitation (averages shown; n=4; error bars, SEM) at the top. The limited DNA cleavage in lane 3 likely results from residual nuclease activity of hEXO1(D173A) that becomes apparent at high protein concentrations (100 nM) in the presence of manganese. **c**, Representative electrophoretic mobility shift assays with hEXO1(D173A)(100 nM) and oligonucleotide based HJ or dsDNA as substrates. Asterisk (*) indicates the position of the radioactive label. The binding buffer contained EDTA (3 mM) and no ATP. No stable DNA binding was detected under our conditions. **d**, Quantitation of electrophoretic mobility shift assays with hMLH1-hMLH3, hMSH4-hMSH5 and hEXO1(D173A), as indicated. The protein-DNA species were resolved in 1% agarose gels. Averages shown; error bars, SEM; n=5. hEXO1(D173A) did not notably affect DNA binding of hMLH1-hMLH3 and hMSH4-hMSH5. **e**, Nuclease reactions with hMLH1-hMLH3 (100 nM), hMSH4-hMSH5 (100 nM) and hEXO1(D173A) (100 nM), carried out at 37 °C in the presence of 5 mM manganese acetate and 2 mM ATP. Linearized DNA (lane 3) was used as a marker.

**Extended Data Figure 7.**
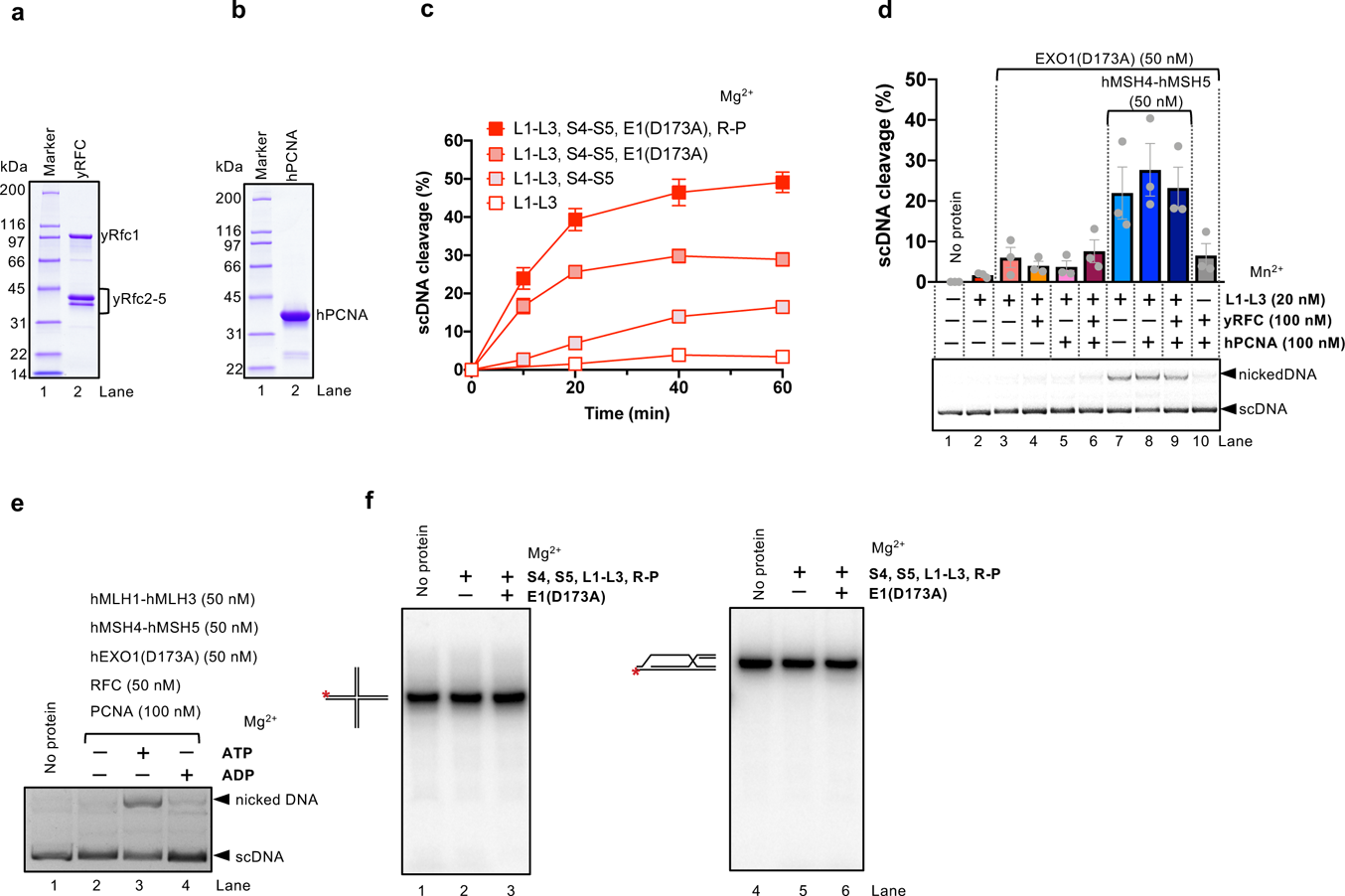
RFC-PCNA do not promote the MLH1-hMLH3 nuclease in reactions with manganese. **a**, Recombinant yeast RFC (yRFC) used in this study. **b**, Recombinant human PCNA (hPCNA) used in this study. **c**, Kinetic experiment carried out with hMLH1-hMLH3, L1-L3 (50 nM); hMSH4-hMSH5, S4-S5 (50 nM), EXO1(D173A) (50 nM) and RFC-PCNA (50-100 nM, respectively), as indicated. Reactions were carried out with 5 mM magnesium acetate and 2 mM ATP at 37 °C. Averages shown; error bars, SEM, n=4. **d**, Nuclease assays with hMLH1-hMLH3 (L1-L3), hMSH4-hMSH5 (S4-S5), hEXO1(D173A) without or with RFC/PCNA, as indicated. The assays were carried out at 37 °C in the presence of 5 mM manganese acetate and 2 mM ATP. A representative experiment is shown at the bottom, a quantitation (averages shown; n=3; error bars, SEM) at the top. Under these conditions, no stimulation of DNA cleavage by RFC-PCNA was observed. **e**, Nuclease assay with indicated proteins and magnesium, either with no co-factor (lane 2), with ATP (2 mM, lane 3) or ADP (2 mM, lane 4). ATP is strictly required for DNA cleavage by the nuclease ensemble. **f**, Nuclease assays with indicated oligonucleotide-based substrates carried out at 37 °C in the presence of 5 mM manganese acetate and 2 mM ATP. All proteins 30 nM. Asterisk (*) indicates the position of the radioactive label. The reaction products were analyzed on a 15% denaturing polyacrylamide gel. No DNA cleavage was observed.

**Extended Data Figure 8.**
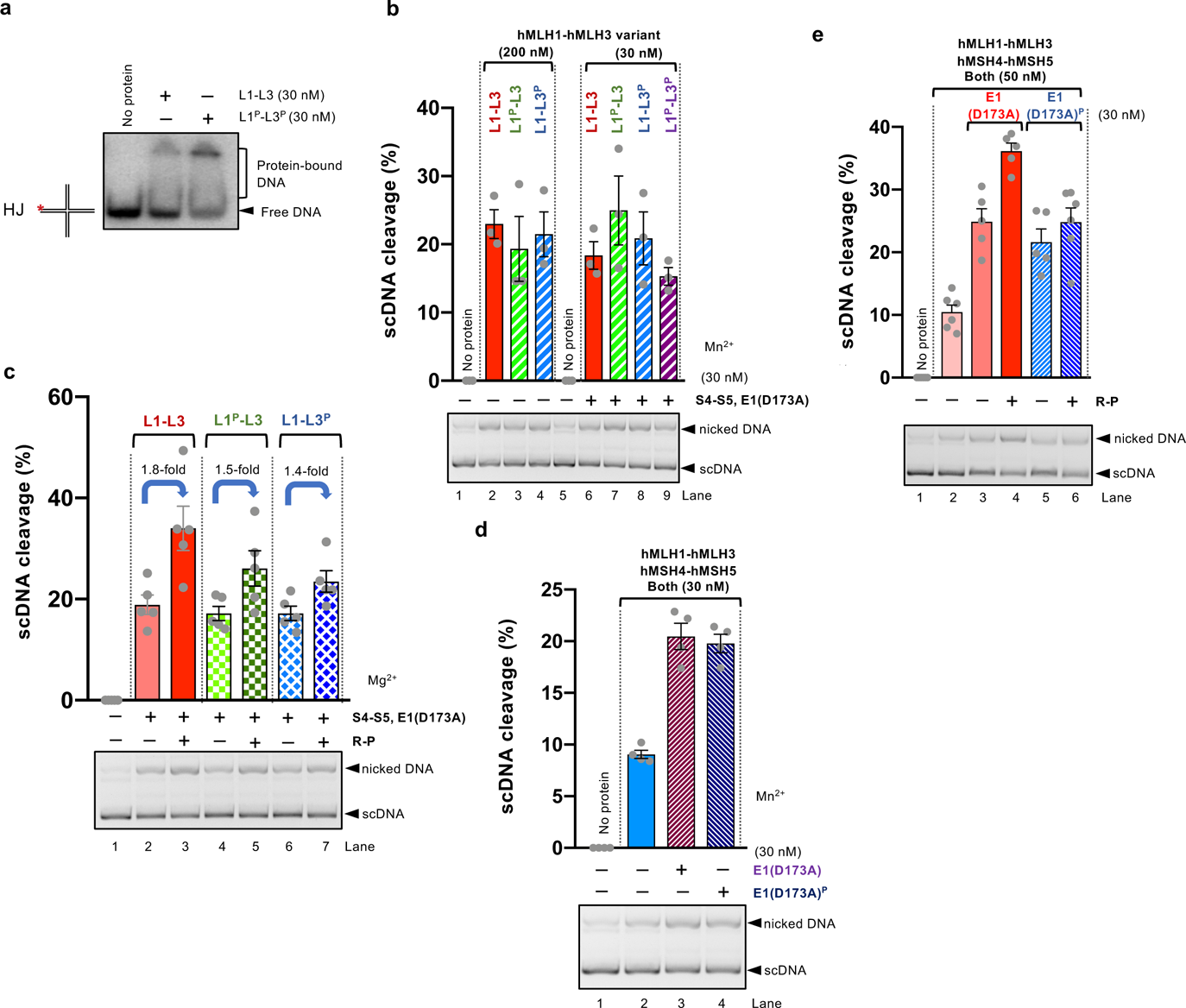
PIP box-like motifs in hEXO1, hMLH3 and hMLH1 facilitate the stimulatory effect of RFC-PCNA on the hMLH3 nuclease ensemble. **a**, The hMLH1^P^-hMLH3^P^ variant (see Fig. 5b) is not impaired in HJ-binding. Electrophoretic mobility shift assay was carried out with 5 ng/μl dsDNA competitor and 3 mM EDTA (no magnesium). **b**, The hMLH1^P^ and hMLH3^P^ variant combinations are not impaired in nuclease activity without or with hMSH4-hMSH5 and hEXO1(D173A) in the absence of RFC-PCNA. The nuclease assays were performed with 5 mM manganese acetate and 2 mM ATP at 37 °C. Averages shown; error bars, SEM, n=3. **c**, Nuclease assays with hMSH4-hMSH5, S4-S5 (50 nM), hEXO1(D173A) (50 nM) and RFC-PCNA (50-100 nM), and a respective hMLH1-MLH3 (L1-L3) variant, as indicated (see Fig. 5b). Mutations in the PIP-box like motif reduce the stimulation of the nuclease ensemble by RFC-PCNA. The assays were carried out with 5 mM magnesium acetate and 2 mM ATP at 37 °C. Averages shown; error bars, SEM, n=5. **d**, The EXO1^P^(D173A) variant with mutated PIP-box motif (see Fig. 5b) is not affected in its ability to promote the nuclease of hMLH1-hMLH3 and hMSH4-hMSH5 (without RFC-PCNA). The assays were carried out with 5 mM manganese acetate and 2 mM ATP at 37 °C. Averages shown; error bars, SEM, n=4. **e**, The EXO1^P^(D173A) variant with mutated PIP-box motif (see Fig. 5b), in complex with hMLH1-hMLH3 and hMSH4-hMSH5 impairs the stimulatory function of RFC-PCNA (50-100 nM). The assays were carried out with 5 mM magnesium acetate and 2 mM ATP at 37 °C. Averages shown; error bars, SEM, n=5.

**Extended Data Figure 9.**
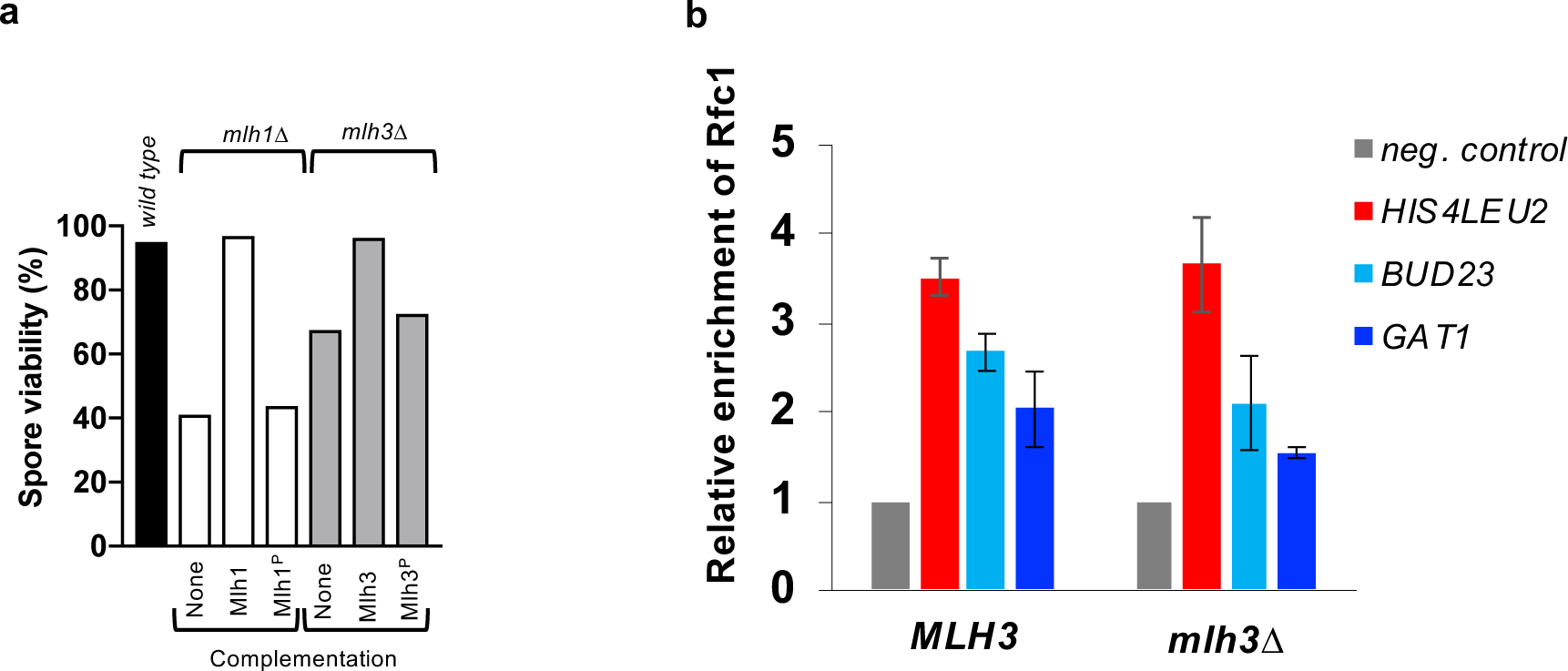
RFC-PCNA regulate meiotic recombination in yeast cells. **a**, Spore viability upon tetrade microdissection, analyzed in the wild type strain, *mlh1*Δ and *mlh3*Δ, and in strains complemented with a construct expressing Mlh1^P^ (Q572A-L575A-F578A) or Mlh3^P^ (Q293A-V296A-F300A). At least 156 spores from 2 biological replicates were analyzed for each genotype. **b**, Rfc1-TAP levels at the three indicated meiotic DSB hotspots relative to a negative control site (*NFT1)* were assessed by ChIP and qPCR in *ndt80Δ* arrested cells after 7 h in meiosis. Mlh3 is not required for the recruitment of RFC to the meiotic DSB hotspots. *MLH3*: VBD2136; *mlh3Δ*: VBD2137. Averages show; error bars, SD, n = 2.

**Extended Data Figure 10.**
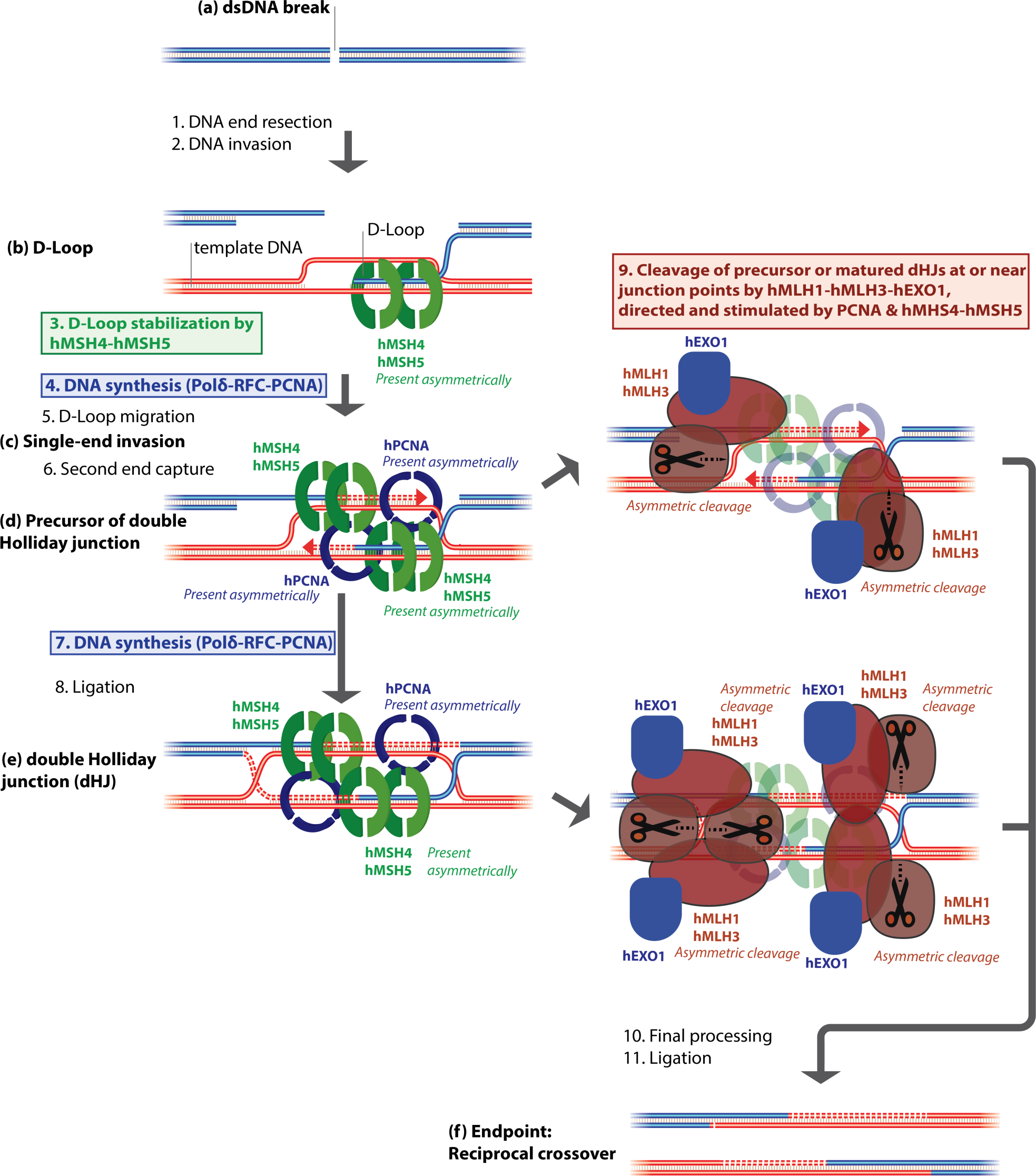
A possible model for biased resolution of recombination intermediates. Meiotic dsDNA breaks (a) are resected (1) and invade matching DNA on a homologous chromosome (2). The unstable D-Loop intermediates (b) are stabilized by hMSH4-hMSH5 (3), DNA synthesis by RFC-PCNA-Polδ (4) and branch migration (5), leading to more stable structures termed single-end invasions (c). This is followed by a second end capture (6), and more DNA synthesis (7) leading to precursors of double Holliday junctions (d) and later matured double Holliday junctions (e). As a result of the previous steps, hMSH4-hMSH5 and RFC-PCNA may be present asymmetrically at the (d) or (e) intermediates at the junctions points or their vicinity. The asymmetric presence of the co-factors then directs and stimulates the biased DNA cleavage (9) of (d) or (e) structures by hMLH1-hMLH3-hEXO1. Upon final processing (10) and ligation (11), the ultimate result is a DNA crossover characterized by reciprocal exchange of the DNA arms of the recombining chromosomes.

